# Three-row stereocilia model predicts mammalian hair bundle behavior

**DOI:** 10.1101/2025.04.17.649156

**Authors:** Varun Goyal, Karl Grosh

## Abstract

Mammalian outer hair cells (OHCs) enhance sound amplification and frequency tuning through stereociliary hair bundles (HBs), which convert mechanical motion into electrical signals via mechano-electric transducer (MET) channels. Experiments show that the HB displacement creeps, and the MET current evinces dual timescales of adaptation in response to mechanical stimulus. Understanding these mechanisms is crucial for elucidating normal auditory function and disorders, yet their origins remain unclear. To address this, we developed a mathematical model of the OHC HB that incorporates three rows of stereocilia with distinct nonlinear adaptive gating mechanisms, nonlinear kinematics, and viscoelastic mechanics. Our model accurately replicates experimental responses to fluid-jet stimulation, predicting simultaneous mechanical creep and slow adaptation of the MET current. Using stiff probe stimulation, the same model reveals even faster adaptation, aligning with experimental observations and emphasizing the stimulus-dependence of the response. The model provides new insights into the functional importance of the three-row stereocilia configuration, offering a mechanistic explanation for its ubiquity in mammalian HBs and its role in facilitating the complex timescales of adaptation.

## Introduction

The ear relies on intricate structures within the cochlea to transduce mechanical stimuli into electrical signals for neural processing. Key to this transduction mechanism is the organ of Corti (OoC), bounded by basilar membrane and tectorial membrane (TM). The OoC comprises various structural-support cells and mechanoreceptors known as outer hair cells (OHCs) and inner hair cells (IHCs) [1]. OHCs primarily amplify and tune sound [2, 3], while IHCs serve as sensory transducers [3]. The motion of stereocilia protrusions (collectively forming a hair bundle (HB)) from the apical surfaces of these hair cells initiates sound processing. In this work, we focus on elucidating the mechanical and electrical dynamics of a mammalian OHC HB. A schematic of the OHC HB is shown in Fig. 1a. Arranged in graded heights, these HBs typically feature three rows of stereocilia, with the tallest stereocilia embedded within the overlaying gelatinous TM [4], and about 16-33 columns of the three-row arrangement oriented in a V or W shape [5]. Each taller row is connected to its shorter neighbor by tip links, composed of cadherin-15 and protocadherin-23 proteins [6], and horizontal top connectors [7]. There are transduction channels, which Beurg *et al*. [8] discovered are localized at the tips of shorter stereocilia within their lipid bilayer. Acoustic excitation induces HB motion, modulating the channel conductance via tip links and allowing cations such as calcium and potassium to permeate through these channels, resulting in the mechano-electric transducer (MET) current. The functional importance of the typical three-row arrangement of stereocilia found in most mammalian HBs remains unclear.

**Figure 1:**
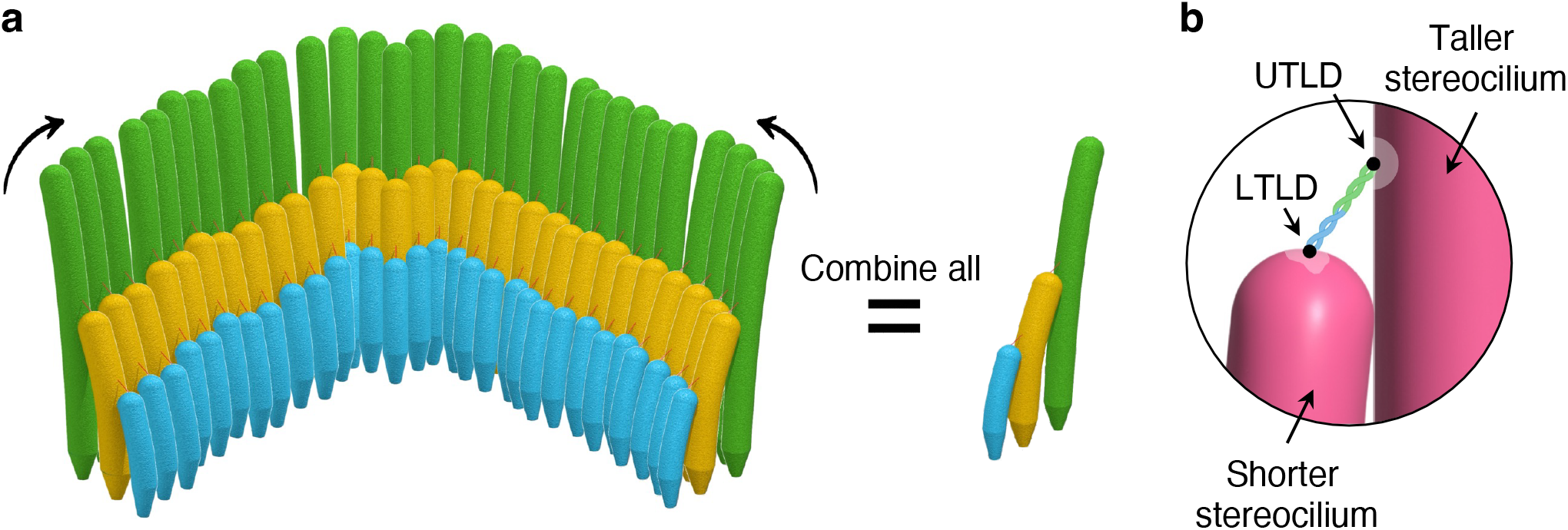
Homogenization of the multi-column OHC HB and definitions of the tip link densities. **a** Simplified representation of a multi-column bundle as a single column. The left side shows a mammalian cochlear OHC HB with 30 columns. Our model reduces this to a single column (shown on the right) to simulate the entire HB’s response. This simplification significantly enhances the model’s computational efficiency, allowing effective incorporation and analysis of nonlinearities while optimizing simulation time (see Methods for details). This approach assumes uniform stimulation of all tallest stereocilia (green) by an external stimulus, necessitating parallel alignment of all columns in contrast to the experimentally observed V or W shape. The HB is not drawn to scale. **b** Two adjacent stereocilia connected by a tip link, illustrating the lower tip link density (LTLD) and the upper tip link density (UTLD).

High-speed video imaging and electrical recordings provide insight into the temporal dynamics of HB displacement and MET current in response to external stimuli [9, 10, 11, 12]. In these experiments, HBs are typically subjected to step inputs delivered via a fluid-jet or a stiff probe, revealing four distinct yet interrelated phases of the electromechanical response: (1) mechanical rise-time; (2) mechanical creep; (3) fast adaptation of the MET current; and (4) slow adaptation of the MET current.

For fluid-jet stimuli with a rise time of ∼0.1 ms [13], HB displacement exhibits an initial rapid ascent (mechanical rise) dictated by the bundle’s viscoelastic properties and the stimulus rise-time. This is followed by a gradual mechanical creep, during which MET currents peak transiently before adapting to a steady-state level. This adaptation process unfolds over a slow time scale (decay constant *>* 10 ms) [12, 14, 15]. In contrast, under rapid and stiff probe excitation, the mechanical rise-time of the HB closely matches the stimulus, typically within tens of microseconds. As a result, HB displacement reaches a steady state without mechanical creep under stiff probe excitation. In contrast, the MET current exhibits distinct fast (0.1–1 ms) [12, 14, 15, 16] and slow adaptation kinetics, differing from those observed with fluid-jet stimulation.

A fifth phase, linked to MET channel activation kinetics, is hypothesized to occur on the order of 1 *µ*s [17]. However, this remains experimentally inaccessible due to the limited bandwidth of patch-clamp recorders [3] and the relatively slow rise-times of conventional piezoelectric stimulators [18]. Consequently, we treat this process as instantaneous in the present model, as suggested by [19]. Understanding the interplay between fast and slow adaptation is critical, as these mechanisms are thought to protect the tip links from excessive strain, preserving HB sensitivity across varying sound levels and mitigating hearing damage from loud stimuli [20, 21].

The molecular composition of stereocilia suggests that calcium binding sites within transduction channels may regulate fast adaptation [16, 22], whereby ion binding increases the force required to maintain equivalent channel open probabilities through channel re-closure [22, 23]. Conversely, slow adaptation may be mediated by the movement of myosin molecules near the upper tip link density (UTLD), where the tip link connects to the taller stereocilium [24], as shown in Fig. 1b. Previous studies [14, 21] discuss differences between these mechanisms, with some groups advocating for fast adaptation being a viscoelastic property of the HB [25, 26].

Theoretical research has focused on developing mathematical models to explain diverse phenomena observed in HBs. The gating-spring model by Howard and Hudspeth revolutionized the field by depicting transduction channels as mechanically-gated ion pathways, opening in response to tension changes in the spring [19, 28]. This foundational model inspired subsequent enhancements [29, 30], including the addition of a “motor” element to replicate slow adaptive characteristics in MET currents [31, 32]. Choe *et al*. [33] introduced a multi-degree-of-freedom kinetic model with dual calcium-binding sites and multiple channel states to simulate fast adaptation dynamics. Subsequent modifications added more channel states and motors [27, 34], increasing model complexity and computational demands. Tinevez *et al*. [25] proposed a simplified two-stereocilia motor model with two degrees of freedom, incorporating active feedback from infiltrating cations. However, as shown in Supplementary Fig. S4, we find that this model can only predict either fast adaptation or slow adaptation, but not both simultaneously, and is thus unable to account for the dual adaptation observed during rigid probe excitation [9, 10, 11, 12]. This limitation arises due to the absence of mechanisms that give rise to a third time constant. Nam *et al*. [35] experimentally demonstrated that uniform stimulation of HBs, influenced by probe shape, resulted in dual adaptive behaviors. While their 3-D HB model supports the observation regarding probe shape dependence, it is indiscernible whether the model can simultaneously predict the two adaptation time constants. Moreover, the model is computationally intensive, involving ∼6300 degrees of freedom and added complexity from stochastic elements. Therefore, we sought to develop a simpler, physiologically inspired three-row model that can accurately replicate the HB’s response to various stimuli, facilitating an intricate understanding of underlying mechanisms.

## Results

### Three-row micro-mechanical model of a mammalian HB

We model the entire multi-column HB as a single representative column, as illustrated in Fig. 1a. This approach is similar to previously established models [19, 28, 25, 33], with the key distinction that our model includes three rows and two MET channels in columnar homogenization. This simplification employs a single Boltzmann probability function for MET channels at the tips of the two shorter stereocilia [8] to represent the fraction of open channels for the entire HB, significantly enhancing computational efficiency compared to multi-column simulations. Within this single column, the three stereocilia are modeled as rigid rods arranged by height, designated as rows 1, 2, and 3, from tallest to shortest, as shown in Fig. 2a. Rotational damping coefficients *λ*_1_, *λ*_2_, and *λ*_3_ model the fluid’s viscous effects, while torsional springs 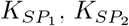 and 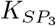 represent rootlet stiffness. Following the gating-spring theory [19, 28], our model incorporates two gating springs 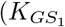 and 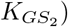 positioned at the tips of rows 2 and 3, at the lower tip link density (LTLD) shown in Fig. 1b. Each UTLD functions as a motile element capable of slipping or climbing [31, 32, 36], regulated by a distinct adaptation complex, or “motor,” which depends on an extent spring 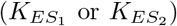 and intracellular viscous coupling (*λ*_*a*1_ or *λ*_*a*2_). The middle row UTLD, being close to its LTLD, is regulated by instantaneous calcium feedback *S* [25], while row 1 UTLD, far from any MET channel, is modeled as calcium-independent. We impose sliding contact constraints between adjacent stereocilia to reflect the influence of horizontal top connectors [7], ensuring the concurrent motion of rows 2 and 3 as the tallest row rotates. The single-column HB homogenization simplifies our model by allowing us to assume that the viscoelastic parameters are aggregated across all columns (30 in our model) and that the channel open probability reflects the density of open MET channels, as shown in equation (1). The three degrees of freedom are bundle rotation *ϕ*, the motion of the first adaptation motor *l*_*a*1_, and the displacement of the second adaptation motor *l*_*a*2_, as illustrated in Fig. 2a, b.

**Figure 2:**
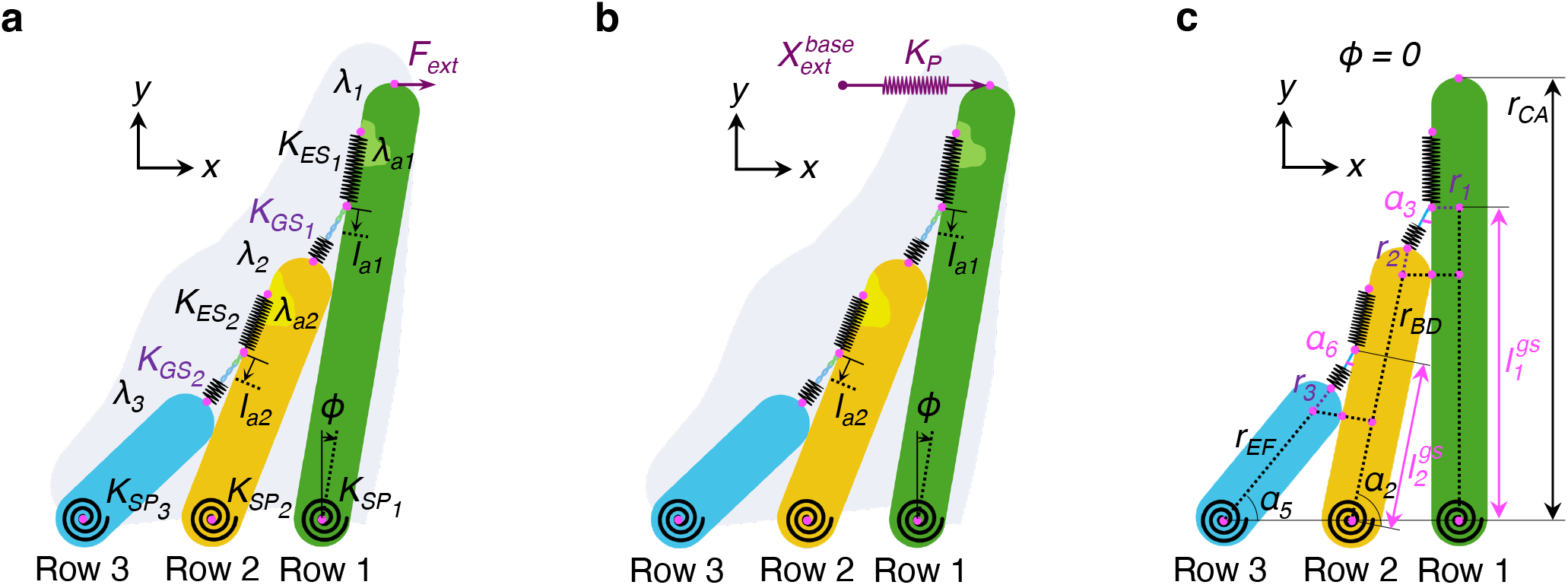
Schematic of the three-row model under fluid-jet (forced) and stiff probe (displacement) excitation. **a** An illustration of the three-row model including viscoelastic components 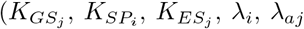 for *i* ∈ [1, 3] and *j* ∈ [1, 2]), as defined in Table 1. The model has three degrees of freedom: HB rotation (*ϕ*) and the motions of the two adaptation complexes (*l*_*a*1_ and *l*_*a*2_). The measurable outputs in response to external force (*F*_*ext*_) are HB displacement (*X*_*hb*_) and MET current (*I*_*met*_). **b** The same three-row model under stiff probe excitation. The probe is depicted as a spring with stiffness *K*_*P*_, driven by a displacement input at its free end. The force experienced by the bundle is 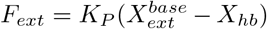. **c** Definition of key geometric lengths and angles relevant to the model when *ϕ* = *l*_*a*1_ = *l*_*a*2_ = 0. Geometric parameter values, derived from Beurg *et al*. [27], are detailed in Table 1. Note that the schematic is not drawn to scale.

The mechanical depiction of the gating mechanism in our model is illustrated in Supplementary Fig. S3. The open probability of a gate at each LTLD (*P*_*o*1_ or *P*_*o*2_) is defined by a two-state Boltzmann relation. This relation is a function of the difference in energies between the open and closed MET channel states, given by

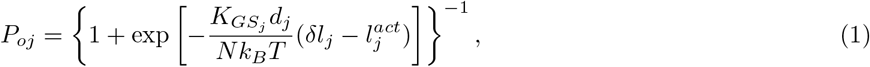

where *j* = 1 and *j* = 2 correspond to LTLDs of rows 2 and 3, respectively. Here, 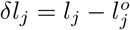 represents the change in length of a gating spring from its natural length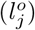, varying nonlinearly with *ϕ, l*_*a*1_, and *l*_*a*2_. The value of 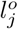is computed when the bundle is upright, as shown in Fig. 2c. The mathematical expressions for *l*_1_ and *l*_2_ are provided in Supplementary Sections 1A-B. Definitions for the remaining parameters are in Table 1.

**Table 1:**
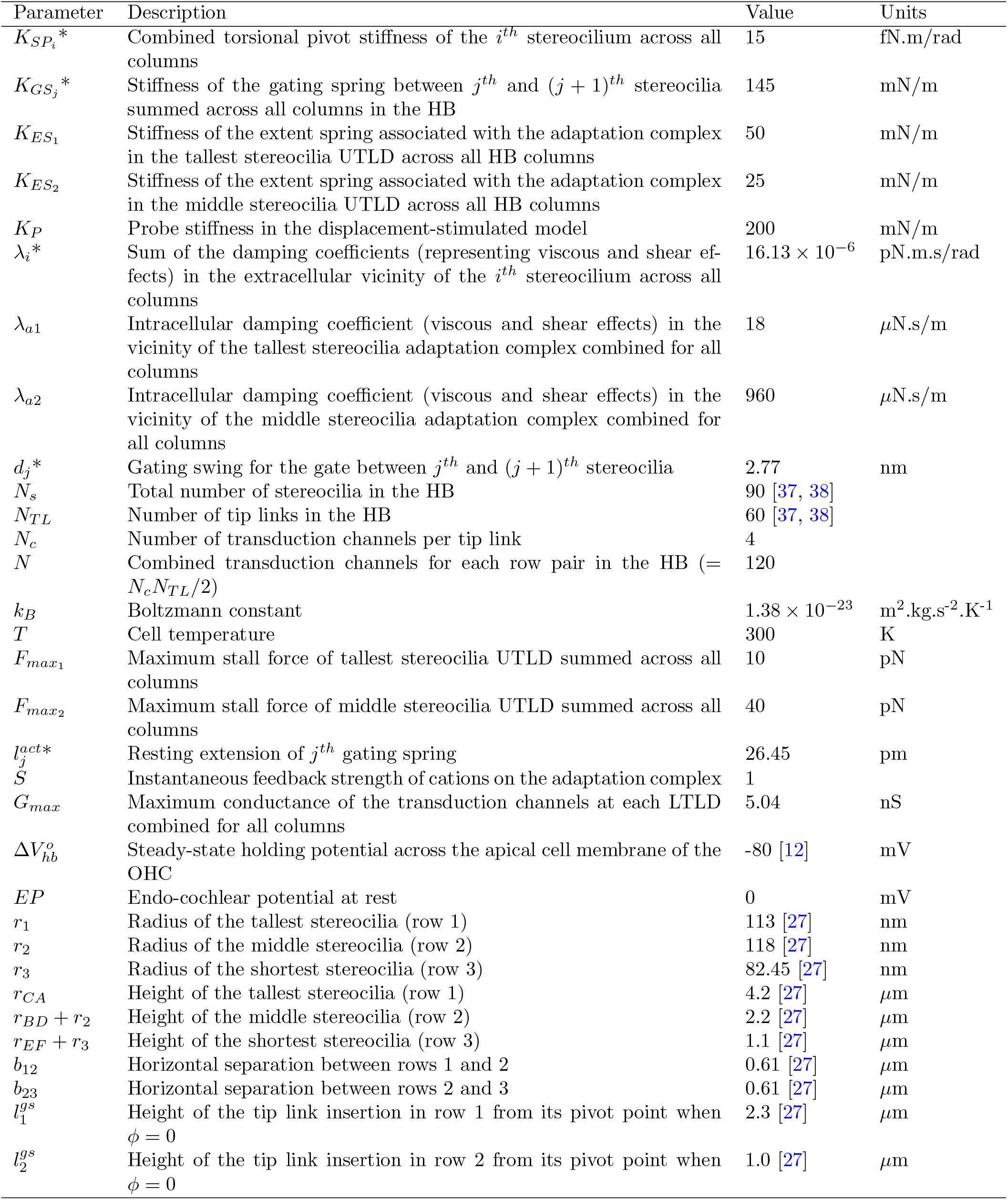
Model parameters, their description, and values. The specified values correspond to the properties of a rat’s OHC HB tuned to approximately 4 kHz. * *i* ∈ [1, 3] and *j* ∈ [1, 2]. Both *i* and *j* are integers.

We derive the nonlinear geometric relationships (detailed in Supplementary Sections 1A-B) and define the energies associated with changes in the HB configuration for each mechanical element (outlined in Supplementary Sections 1E-F). Using Lagrange’s equations, we obtain three coupled equations of motion incorporating both geometric and constitutive nonlinearity from channel gating. These equations govern HB dynamics in response to external inputs, such as a force *F*_*ext*_(*t*) (equation (9)) from a fluid-jet as illustrated in Fig. 2a, or a displacement 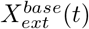 (equation (10)) from a probe, as shown in Fig. 2b. In the ear, this represents excitation from the TM moving the tallest stereocilia. The dynamic equations for HB stimulated by a force are:

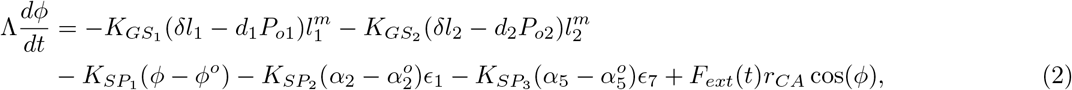

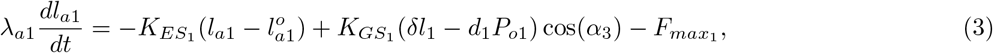

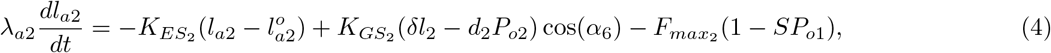

where 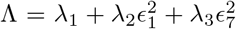 is the equivalent extracellular damping coefficient and the quantities *ϵ*_1_ and *ϵ*_7_, which depend on *ϕ* are defined in Supplementary Table S2. The angles *α*_2_, *α*_3_, *α*_5_, and *α*_6_ (shown in Fig. 2c) also depend on *ϕ*, with *α*_3_ and *α*_6_ additionally dependent on *l*_*a*1_ and *l*_*a*2_, respectively. 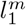 and 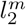 represent the moment-arm lengths displayed in Supplementary Fig. S2b, with mathematical expressions in Supplementary Section 1F. Constants with superscript “*o*” are evaluated when *ϕ* = *l*_*a*1_ = *l*_*a*2_ = 0, and remaining model parameters are listed in Table 1. The relation between the model outputs *ϕ, l*_*a*1_, *l*_*a*2_ and experimental displacement *X*_*hb*_ and MET current *I*_*met*_ is given by

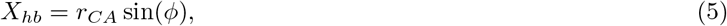

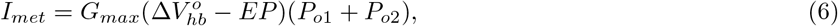

where the geometric and electrical properties are defined in Table 1. For a displacement input from a stiff probe with stiffness *K*_*P*_, *F*_*ext*_(*t*) in equation (2) changes to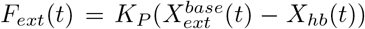, resulting in the following modification:

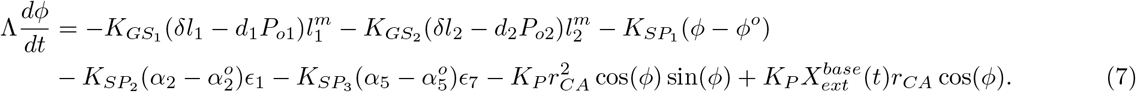

The response of the HB is governed by equations (7), (3), and (4). Equation (7) shows that this system is both stiffened and forced by *K*_*P*_ proportional terms.

Three key equations describe the bundle dynamics in our model. The bundle’s rotation, as outlined in equation (2) or (7), is influenced by torque from two gating spring forces 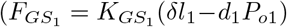 and 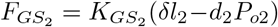, shown in Supplementary Fig. S2b), resistance from torsional springs at the three pivots, and equivalent fluid damping (Λ). For a displacement input, equation (7) shows that the probe’s stiffness further constrains bundle rotation in the excitatory direction, effectively stiffening the HB system. Equations (3) and (4) describe the dynamics of the two adaptation motors. The row 1 motor is driven by the component of 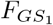 along the extent spring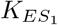, given by 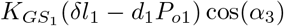 in equation (3). Similarly, the row 2 motor is driven by the component of 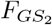 along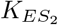, given by 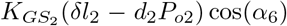 in equation (4). The last terms of equations (3) and (4) represent the effect of tensioning in the model. Given the ample evidence supporting the calcium dependence of slow adaptation [10, 12, 40], we incorporate this mechanism into the middle row UTLD through an active feedback term given by 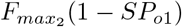 in equation (4). However, the calcium dependence of fast adaptation is a subject of debate in the literature [39, 40]. In our model, we posit that fast adaptation is calcium-independent and governed by the row 1 UTLD, as represented in equation (3) by a passive tensioning force,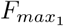. A detailed derivation of our model can be found in Supplementary Section 1.

### Three-row model quantitatively matches creep and slow adaptation

To analyze creep and slow adaptation, we stimulated the HB with step forces of varying amplitudes, each with a rise-time of 0.1 ms, similar to the experimental fluid-jet forcing used by Caprara *et al*. [12], using a single set of model parameters (Table 1). The model results are compared with experimental data in Fig. 3a, b. As shown in Fig. 3a, our model provided a quantitative fit of the mechanical response, reproducing the mechanical rise and subsequent creep of the HB displacements for all forces. Figure 3b compares the MET currents, demonstrating that the model quantitatively fits the slowly adapting currents up to ∼70 nm HB displacement. However, the model predicted a more rapid current saturation than observed experimentally, possibly due to the homogenization of the multi-column HB into a single column.

**Figure 3:**
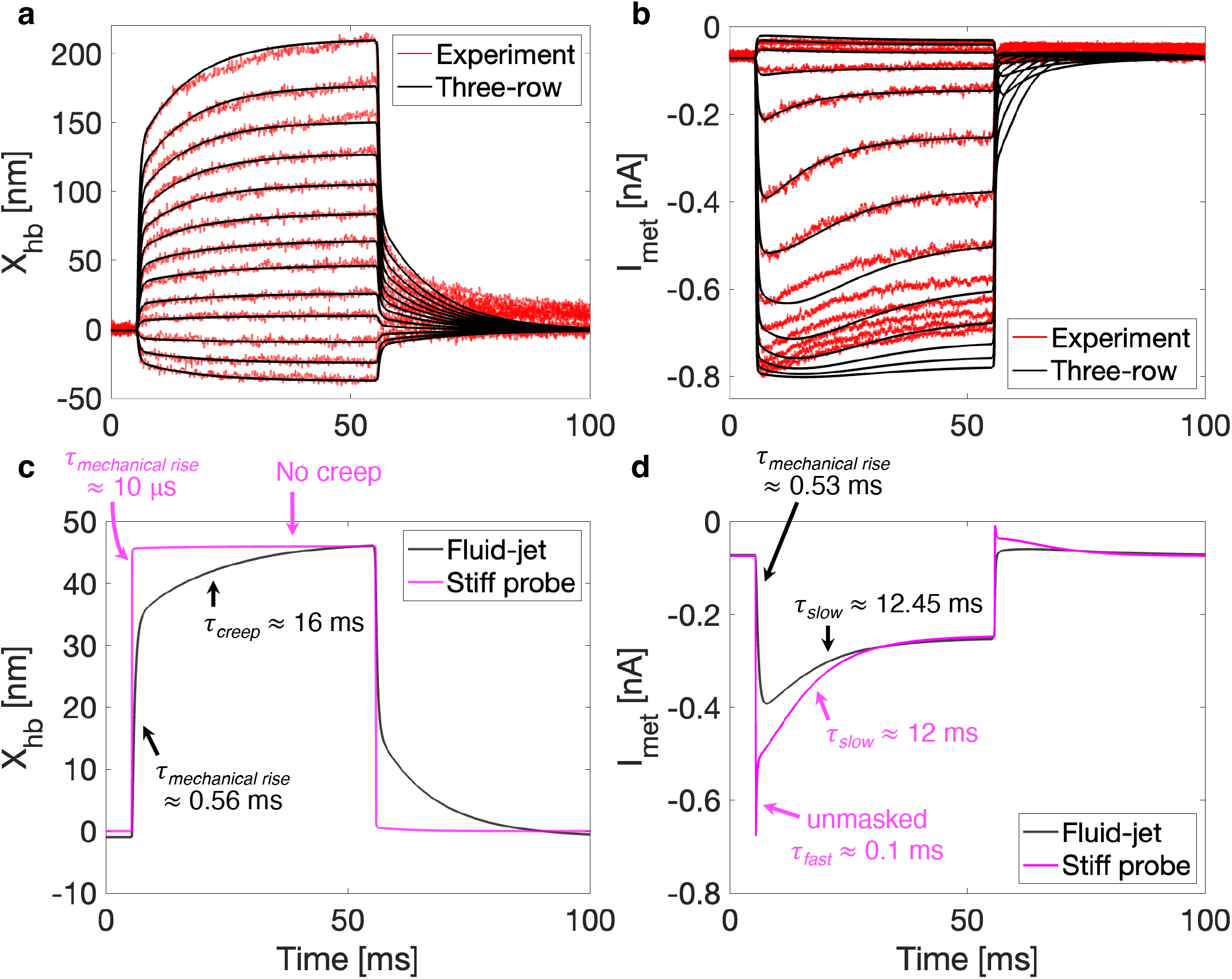
Three-row model quantitatively predicts the fluid-jet experiments and three response time constants. **a** HB displacements in response to thirteen distinct force amplitudes. **b** Corresponding MET currents for the same forces. The forces replicate a fluid-jet with a 0.1 ms rise-time and amplitudes of -180, -115, -41, 51, 127, 217, 291, 375, 470, 569, 677, 797, and 950 pN. The model illustrates an initial mechanical rise followed by a creep in displacement, with currents showing a rapid rise and slow adaptation, consistent with experimental data (red). **c** Comparison of HB displacement responses between a stiff probe (magenta) and fluid-jet (black) model. **d** Comparison of associated MET currents from both models. The probe had a 10 *µ*s rise-time and a 47 nm displacement amplitude, and the fluid-jet data corresponds to 217 pN force amplitude. Probe stimulation resulted in a faster mechanical rise (∼10 *µ*s) due to HB stiffening, resulting in no creep, a higher MET current peak, and revealing fast adaptation (∼0.1 ms), absent in fluid-jet stimulation. Slow adaptation followed with a decay constant of ∼12 ms. The estimates of mechanical rise, fast, and slow adaptation were obtained by fitting the magenta curves in **c** and **d**. Fitting the fluid-jet response (black) suggested a similar mechanical rise time for both displacement (∼0.56 ms) and current (∼0.53 ms), along with the creep (∼16 ms) and slow adaptation (∼12.45 ms) time constants.

### The same three-row model reveals fast adaptation during rigid probe excitation

We stimulated the HB model using a stiff probe, as illustrated in Fig. 2b. In this setup, the base of the spring, controlled experimentally by a piezo actuator, was displaced with a rise-time of 10 *µ*s. Aside from the added spring stiffness from the probe, the HB parameters were identical in both fluid-jet and probe simulations. The results from the probe model are presented in Fig. 3c, d. Figure 3c compares HB displacement from the probe model (magenta) to the fluid-jet model (black) for similar bundle displacements. The added stiffness from the probe reduced mechanical rise-time, eliminated creep, and revealed fast adaptation. Due to the faster mechanical activation, the MET current peak was nearly 1.7 times higher than that in the fluid-jet case, as shown in Fig. 3d. The MET currents showed both fast and slow adaptation, consistent with experiments [9, 10, 11, 12]. Our analysis suggests that fast adaptation was obscured due to the slower mechanical rise in fluid-jet experiments and the lower stiffness of such a setup.

We compared our three-row model with the two-row model developed by Tinevez *et al*. [25] under both fluid-jet and stiff probe stimulations. We optimized the parameters of the two-row model to fit fluid-jet experimental data [12] (Supplementary Table S3) and applied the same optimized model to the probe stimulation, incorporating probe stiffness. As demonstrated in Supplementary Fig. S4, the two-row model can generate creep and slow adaptation under fluid-jet stimulation. However, it cannot predict both fast and slow adaptation even with added probe stiffness due to the absence of mechanisms for a third time constant. Fast adaptation is essential for transducing high frequencies typical in mammalian hearing, enabling precise cycle-by-cycle amplification [41], whereas slow adaptation possibly plays a protective role by preventing cellular damage. The three-row model accounts for both adaptation mechanisms, offering a mechanistic explanation that the two-row model cannot provide. This underscores the critical role of the third row of stereocilia in facilitating adaptation in the middle row and highlights the significance of incorporating three rows to fully capture the temporal dynamics under varied loading conditions.

### Prediction of HB operating range, sensitivity, and stiffness

We evaluated the accuracy of our nonlinear model in predicting fundamental biophysical quantities, including the bundle’s operating range (OR), sensitivity, and stiffness. The OR, defined as the displacement span where 10% to 90% of transduction channels are open, was estimated from activation curves constructed for both fluid-jet and stiff probe models. Figure 4 compares the fluid-jet model (black) and stiff probe model (magenta) against fluid-jet experimental data (red) [12] and rigid probe data from mice (blue) [10]. The fluid-jet model’s OR of ∼86 nm closely matched the experimental value of ∼77 nm. However, it underestimated sensitivity, defined as the maximum slope, by ∼30%, possibly due to the single-column HB approximation or unaccounted transduction channel elements, explored further in the Discussion. The probe model predicted an OR of ∼40 nm, consistent with experimental values in mice [10, 11] while underestimating sensitivity by ∼10%.

**Figure 4:**
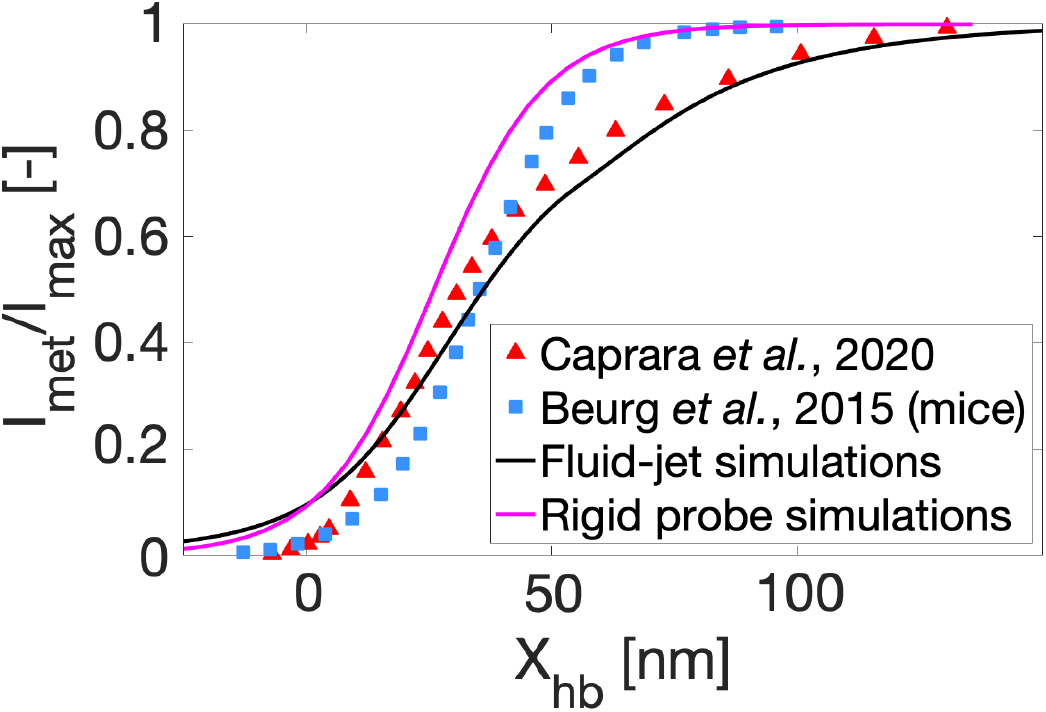
Predicted HB operating range (OR) consistent with experiments. Activation curves derived from normalized peak currents and corresponding HB displacements for each applied force or displacement amplitude provide a measure of the HB’s OR (10-90% width). The fluid-jet experimental OR is approximately 77 nm (red triangles), while the fluid-jet model predicts an OR of ∼86 nm (black solid line). The OR from the stiff probe model is estimated at 40 nm (magenta solid line), aligning with experimental measurements in mice from Beurg *et al*. [10] (blue squares).

Using our model, we also assessed bundle stiffness (*K*_*HB*_) and compared it to experimental observations under two conditions: intact and severed tip links. Disruption of tip links has been experimentally shown to decrease *K*_*HB*_, indicating the contribution of gating springs [15, 27, 37]. We derived the force-displacement relation for both conditions to estimate *K*_*HB*_ from its slope, as described in Supplementary Section 1H. For the intact tip links model, we used our nonlinear three-row model. We eliminated all gating springs for the disrupted tip links model by setting their stiffness 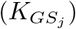 to zero, thereby deactivating transduction and adaptation mechanisms. Our stiffness estimates from the fluid-jet and probe models for the two conditions were in agreement with each other. Using the intact tip links model, we determined *K*_*HB*_ = 4.5 mN/m when ∼40% of MET channels were open, aligning with experimental values for apical HBs in rats [37]. In the disrupted tip links model, *K*_*HB*_ was estimated at 2.34 mN/m, a 48% reduction from the intact model, suggesting that gating springs contributed ∼48% to bundle stiffness. Our results are in reasonable agreement with the observed 38% decrease in bundle stiffness in rats [27, 37] and mice [15] after tip link disruption.

### Geometric gain requires precise definition

We analyzed the “geometric gain,” which measures the projection of gating spring displacement onto the horizontal displacement of the HB. Traditionally, the geometric gain is approximated as the ratio of stereociliary spacing (*b*_*st*_) to the tallest row’s height (*h*_*st*_) [29], based on three assumptions: infinitesimally thin stereocilia, gain as the ratio of UTLD slip to the tallest stereocilium’s horizontal motion, and small HB rotations. However, an accurate measure should account for the change in gating spring length and angular orientation rather than just the slip at the contact point. Our model provided a means to estimate this value precisely.

We calculated two geometric gains, *γ*_1_ and *γ*_2_, for the two gating springs as time-dependent functions in Supplementary Section 1I. The exact initial values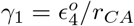 and 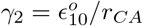 were 0.11 and 0.22, respectively (*ϵ*_4_ and *ϵ*_10_ defined in Supplementary Table S2). Using geometric parameters from Table 1, *b*_*st*_*/h*_*st*_ was 0.14, 27% higher than the 0.11 predicted by our model for an apical OHC HB of a rat.

We further compared *b*_*st*_*/h*_*st*_ and 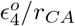 at five cochlear locations (30 kHz, 12 kHz, 8 kHz, 2 kHz, and 750 Hz) using morphometric data from Yarin *et al*. [5] for adult guinea pigs, assuming a constant *b*_*st*_ = 0.5 *µ*m [41, 43], as detailed in Supplementary Fig. S5. The *b*_*st*_*/h*_*st*_ ratios were 0.25, 0.17, 0.15, 0.11, and 0.08 from high- to low-frequency locations, while 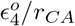 predictions were 0.123, 0.098, 0.083, 0.06, and 0.052, respectively. This comparison indicates that *b*_*st*_*/h*_*st*_ consistently overestimates geometric gain by a factor of 1.5 to 2, indicating that models and experiments relying on this ratio likely misestimate variations in gating spring length, tension, bundle sensitivity, and stiffness.

While *γ*_1_ and *γ*_2_ vary with time and the three degrees of freedom, their dependence on time and stimulus level remains minimal. Even for large HB displacements (∼200 nm), well beyond physiological ranges, the gains increase by only 4%. This justifies the assumption of a constant gain, consistent with previous studies [29, 44]. However, estimating geometric gain solely as *b*_*st*_*/h*_*st*_ is imprecise, as it neglects the influence of stereocilia morphology, particularly radii, and the dynamic reorientation of tip links. Crucially, variations in tip-link length and angle (*α*_3_ and *α*_6_ in Fig. 2c) play a significant role and must be accounted for in gain estimation.

### Mapping response conditions using a linearized approach

The linearized analysis enables closed-form computation of small-signal time constants and facilitates exploring these diverse response characteristics. We derived a linear three-row model by linearizing the nonlinear system (equations (1)-(7)) about an equilibrium state corresponding to the experimental resting open probability (∼7%), as detailed in Supplementary Section 2. This yielded a third-order Jacobian matrix whose eigenvalues and eigenvectors define the system’s mechanical and electrical response to small perturbations (*ϕ* ≪ 1). The resulting closed-form solution is given by

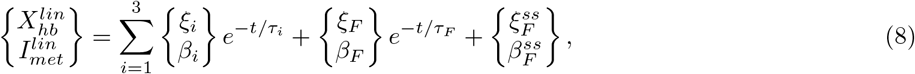

where *τ*_1_, *τ*_2_, and *τ*_3_ are related to system eigenvalues representing mechanical rise, fast, and slow adaptation time constants, respectively. The coefficients *ξ*_*i*_ and *β*_*i*_ are determined by the system eigenvectors and initial conditions (Supplementary Section 2E), and the final two terms capture the response to a force with finite rise-time *τ*_*F*_.

Under fluid-jet stimulation, linearization using the parameters in Table 1 produced characteristic time constants of *τ*_1_ = 0.11 ms, *τ*_2_ = 0.43 ms, and *τ*_3_ = 12.82 ms. This linearized model accurately predicted bundle displacements up to 50 nm but consistently underestimated the MET currents, as shown in Supplementary Fig. S6. To investigate the origins of creep and slow adaptation, we compared normalized experimental displacement and MET current responses to their theoretical counterparts (equation (8)). As shown in Fig. 5, experimental data (red) and theoretical solutions (black) revealed that the linearized system qualitatively captured the overall response trends. We found that the creep emerged when the ratios *ξ*_2_*/ξ*_1_ and *ξ*_3_*/ξ*_1_ were positive (Fig. 3a), consistent with the multi-timescale relaxation in displacement. In contrast, slow adaptation was associated with a positive *β*_2_*/β*_1_ and a negative *β*_3_*/β*_1_, with the magnitude of the latter governing the extent of adaptation (Fig. 3b).

**Figure 5:**
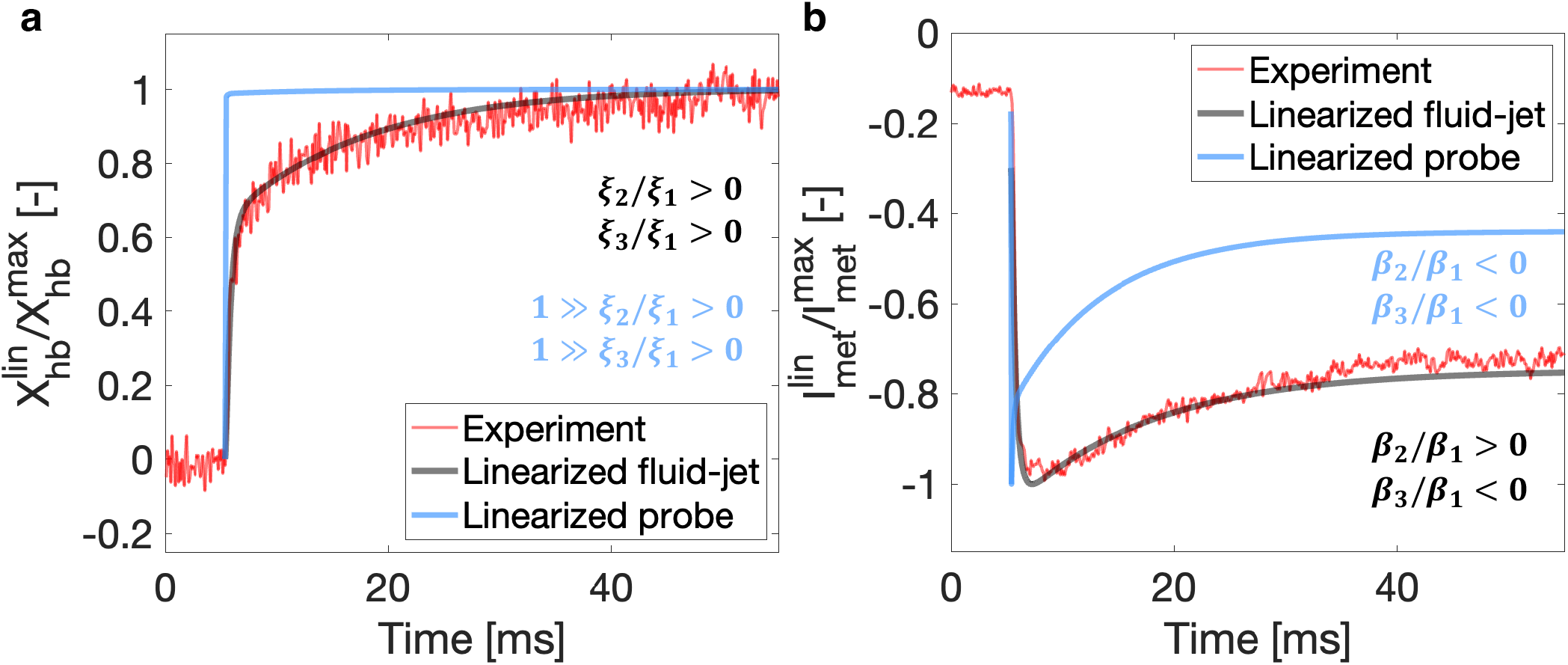
Linearization of the three-row model reveals how response coefficients govern mechanical creep, slow adaptation, and fast adaptation. **a** Normalized HB displacement trace from the linearized fluid-jet model (black) compared with experimental data (red), each normalized to its peak absolute value. The linearization, using parameters from Table 1, accurately represents experimental trends. The polarity of the response coefficient ratios (*ξ*_*i*_, *i* ∈ [1, 3]) dictates the direction and extent of mechanical creep. The response of the linearized probe model (blue) shows the same polarity for both ratios, but their small magnitudes render *ξ*_1_ and *τ*_1_ dominate the response. **b** Normalized MET current trace from the linearized model (black) compared with experimental data (red). The model underestimates the peak current (see Supplementary Fig. S6), leading to an elevated normalized resting current (initial offset of the black trace). The polarity of the MET response coefficients (*β*_*i*_) governs slow adaptation in the fluid-jet model. Incorporating probe stiffness shortens the linearized time constants and reverses the polarity of *β*_2_*/β*_1_, revealing fast adaptation (blue).

When the linearized system was stimulated using a stiff probe, the effective time constants were decreased to *τ*_1_ = 13 *µ*s, *τ*_2_ = 0.14 ms, and *τ*_3_ = 8.29 ms. The probe’s added stiffness resulted in a displacement that was dominated by the fastest exponential mode, *ξ*_1_ exp(−*t/τ*_1_), because the participation ratios *ξ*_2_*/ξ*_1_ and *ξ*_3_*/ξ*_1_ were small (see Fig. 3c). Creep was suppressed in the stiffened system. Under stiff probe stimulation, the polarity of *β*_2_*/β*_1_ was reversed, triggering fast adaptation that was absent in the fluid-jet case. The amplitudes of fast and slow adaptation were modulated by |*β*_2_*/β*_1_| and |*β*_3_*/β*_1_|, respectively, both of which scaled inversely with the elastic stiffness 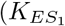 or 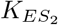 in Table 1). Moreover, when *τ*_*F*_ *> τ*_2_ *> τ*_1_, the slow onset of force masks the fast adaptation dynamics, as previously described [42]. Therefore, under the parameter values used in our model (refer Table 1), rapid stimulation and stiffening of the HB are necessary to observe fast adaptation.

The linearized analysis reveals three key insights. First, the mechanical and electrical behavior is governed by the same three time constants (*τ*_*i*_). Second, differential weighting of the response coefficients underlies the mechanistic basis of creep, fast adaptation, and slow adaptation. Third, the presence of the third stereociliary row is critical for adaptation in the middle row, as it introduces an additional mechanical degree of freedom (*l*_*a*2_) and gives rise to the third time constant.

To address the underestimation of MET currents by the fully linearized model (Supplementary Fig. S6), we constructed a model that retained the constitutive nonlinearity while simplifying geometric constraints (Supplementary Section 3). Comparative simulations under both fluid-jet and probe stimulation (Supplementary Fig. S7) revealed that this model closely tracked the fully nonlinear model, maintaining mechanical and electrical fidelity up to ∼75 nm of HB displacement. This strong agreement across a broad displacement spectrum supports two observations: (1) the validity of a time-independent geometric gain, an assumption adopted in prior models [29, 44] and retained in the partially linearized three-row framework, and (2) that the three-row model, with only constitutive nonlinearity, provides a reliable approximation within the physiological range of HB displacements, without substantially underestimating MET currents as observed in the fully linearized case.

## Discussion

In this study, we developed a simplified three-degree-of-freedom model representing the dynamics of the multi-column HB. This model reproduces four characteristic response phases (mechanical activation, creep, and dual adaptation) observed in experimental recordings. The separation of fast and slow adaptation dynamics emerges naturally from two viscoelastic adaptation mechanisms, each arising from the structural asymmetry introduced by the three-row configuration. The spatial localization of MET channels at the tips of rows 2 and 3 [8] suggests that row 1 UTLD, being more distal from all MET channels, likely operates independently of calcium influx, consistent with the observed calcium-independent fast current decay [39, 45]. Consequently, under our model assumptions, the fast decay of the current [39, 45] is attributed to the intrinsic viscoelastic response of the row 1 UTLD 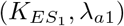.In contrast, the calcium-dependent slow adaptation [10, 12] is governed by row 2 UTLD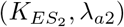, located in close proximity to the MET channels in the middle row. These findings underscore the potential role of the three-row architecture, as the absence of the third row would decouple row 2 UTLD from the system, abolishing the integrated mechanoelectrical behavior seen in experiments.

We further demonstrated that the observability of fast adaptation is contingent on both stimulus type and its rise-time. Simulations of our model under fluid-jet and stiff probe stimulation revealed marked differences in mechanical and electrical responses (Fig. 3). During probe stimulation, displacement creep was eliminated, and MET currents exhibited fast adaptation, consistent with experiments [9, 10, 11, 12]. This finding is important *in vivo*, where the TM attachment stiffens the OHC HB [43]. For a guinea pig, we estimated the TM shear stiffness at ∼7.8 mN/m for the 4 kHz location based on data from Richter *et al*. [46]. This level of TM stiffness is sufficient to stiffen the HB and unmask fast adaptation. Additionally, the presence of the other two OHC HBs in the cross-section may further stiffen the system, enhancing sensitivity and enabling rapid adaptation to prevent excessive tension in the tip links during loud sound exposure.

The dynamic three-stereocilia model offers significant flexibility in integrating mechanisms at both LTLDs and UTLDs. Recent advances in understanding the multi-protein composition of MET channels have identified key proteins such as TMIE (transmembrane inner ear), TMC1/TMC2 (transmembrane channel-like) [47], and accessory proteins like LHFPL5 (lipoma HMGIC fusion partner-like 5) [15], CIB2 (Ca^2+^ and integrin-binding protein) [48], and PIP_2_ (phosphoinositol-4,5-bisphosphate) [47]. Beurg *et al*. [15] suggested that at the LTLD, protocadherin-15 anchors the MET channel complex, interacting with TMC1 and LHFPL5, while CIB2 tethers this complex to the actin filamentous structure [49, 50]. Effertz *et al*. [51] found that PIP_2_ depletion affects channel conductance and cation permeability, abolishing fast adaptation and increasing resting open probability in rat cochleae. At the UTLD, cadherin-23 associates with actin filaments through a complex involving harmonin-b, myosin VIIa, and sans, regulating tip link tension and current adaptation [52, 53].

Our three-row model allows the integration of additional viscoelastic elements to generate adaptive or rate-dependent forces at LTLD or UTLD sites, potentially mimicking multi-protein interactions. For instance, proteins such as TMIE, TMC1/TMC2, PIP_2_, and LHFPL5 could be represented by viscoelastic components whose forces dynamically respond to changes in cell holding potentials [12, 54] and calcium concentrations [12, 27]. These modifications may also provide evidence that fast adaptation is a viscoelastic phenomenon [19, 25, 26, 28]. Detailed modeling of these protein interactions would require additional parameters and increased computational complexity. However, our three-row model is well-suited for these mechanistic numerical experiments, holding the potential to offer valuable insights into protein interactions at the molecular level.

HBs *in vivo* function in concert with thousands of other HBs, intricately connected to numerous elements within the OoC. While simplified HB models, like the one in this study, focus on the micro-nano mechanics of isolated HBs, they can be incorporated into comprehensive cochlear models [41, 43]. These global models simulate fluid-structure, electrochemical, and structure-structure coupling to understand sound processing in the inner ear. Our linearized three-row model can help determine whether a partially linear or fully nonlinear model is needed based on TM displacement amplitudes in the global model. A key feature of global models is OHC-driven amplification, where OHC motility is governed by cell depolarization from ion influx through mechanically-gated channels in HBs [3]. Parameters like HB stiffness, sensitivity, and MET channel conductance, derived from HB models, are crucial for refining these predictions. Thus, an accurate physiological representation of HBs, as presented in this study, is indispensable for effectively modeling cochlear dynamics.

## Methods

### Simulations

We use MATLAB to generate and solve geometric symbolic equations. The time-domain solution is obtained with a custom fourth-order Runge-Kutta method. In this study, we stimulate the HB with either a step-like force or a step-like probe displacement. The applied force, *F*_*ext*_(*t*) in equation (9), and the probe’s base displacement, 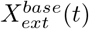 in equation (10), both have a rise-time *τ*_*F*_ and amplitudes *F*_*o*_ and 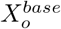,respectively.

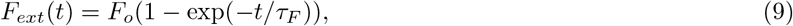

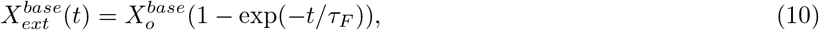

where the subscript “*F* “ denotes force. For displacement input, the subscript “*F* “ is retained for the rise-time because it appears as 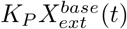 in equation (7), representing a force. The forces were configured to mimic the step-like stimuli from a fluid-jet in experiments by Caprara *et al*. [12], with a rise-time of ∼0.1 ms [13]. We adjusted our model’s force amplitude *F*_*o*_ to match the experimental displacements and MET current responses at 0.1 mM intracellular BAPTA under a −80 mV holding potential [12]. The displacements 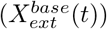 were designed to replicate the action of a stiff probe in the experiments, which had a rise-time of ∼10 *µ*s [12].

Using MATLAB’s parallel computing toolbox, we simultaneously solved the bundle response at various input levels. With 30 workers on the Great Lakes High-Performance Computing cluster, it takes ∼0.5 ms to compute a 1 ms bundle response with a 1 *µ*s time step. On a MacBook Pro with 16 GB RAM and an M1 chip using 8 GPUs, the same computation takes roughly 10 ms. For our simulations, we ran the bundle response for 160 ms per input: 55.4 ms pre-force time, 50.4 ms applied force duration, and the remaining time post-input.

### Mechanical, electrical, and geometric parameters in the model

The model comprises 20 mechanical parameters, 3 electrical parameters, and 13 parameters associated with bundle morphology. This makes individual estimation of all parameters computationally challenging. To streamline this process, we enforce parameter equality among analogous mechanical elements: 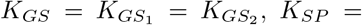 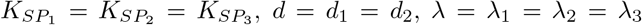, and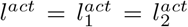. Simulating both fast and slow adaptation dynamics requires 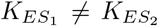 and *λ*_*a*1_ ≠ *λ*_*a*2_ (refer to Supplementary Section 2F and Table 1 for details). To match the resting bundle displacement and MET current with experiments [12], we set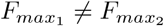. These constraints reduce the number of mechanical parameters to 12. For the electrical parameters, holding potential corresponds to experiments [12], maximum channel conductance in a row-pair *G*_*max*_ was estimated from Johnson *et al*. [55], and *EP* was assumed to be 0 mV. The geometric HB morphology was derived from rat apical HBs [27].

### Determination of mechanical parameters

To set mechanical parameters, we initialized *K*_*GS*_ and *K*_*SP*_ using physiological estimates from Tobin *et al*. [37] for a rat OHC HB. The remaining parameters were either based on previous studies [25, 32, 33, 35, 36] or were assumed. For example, *λ* was derived by observing the mechanical rise of HB displacement in experiments [12]. We then performed a sensitivity analysis on the experimental data and our initialized model, varying single or paired parameters to minimize the error between theoretical predictions and experimental data at each time step. The objective function (mean RMS error) was minimized for both displacement (E_*hb*_) in equation (11) and MET current (E_*met*_) in equation (12).

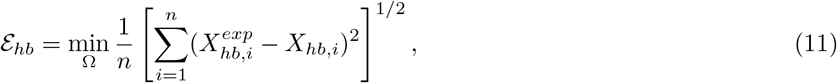

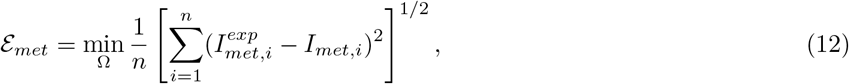

where Ω represents a single or paired parameters, 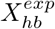 is the experimental bundle displacement, 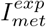is the experimental MET current, and *n* is the number of time steps in the data. Note that for each Ω, the force amplitude (*F*_*o*_) was also iteratively altered to minimize the RMS errors.

To simulate both fast and slow adaptation, we used the linearized three-row model to determine 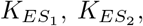, *λ*_*a*1_, and *λ*_*a*2_ (Supplementary Section 2F). We confined *S* between 0 and 1 to prevent sign changes of the motor stall force 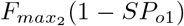.A higher *S* indicates stronger cation feedback, while *S* = 0 eliminates cation dependence. With all parameters determined (Table 1), we used a single parameter set for nonlinear simulations to assess HB response to varying force (*F*_*o*_) and displacement 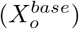 amplitudes.

## Acknowledgements

The authors thank the NIH grant NIH-NIDCD 04084 for providing financial support for this work. Furthermore, the authors extend their gratitude to Dr. Anthony Peng and Dr. Maryline Beurg for generously providing the experimental data, which was instrumental in testing our model.

## Author contributions statement

V.G. and K.G. designed the study, conducted the research, analyzed the data, and authored the manuscript.

## Additional information

### Supplementary information

The nonlinear model developed in this study is described in detail in Supplementary Section 1. The linearized three-row model is covered in Supplementary Section 2, and the model with linearized geometry is derived in Supplementary Section 3.

## Data availability

Simulation data are available upon request or can be generated using our open-source code and visualization tool. The experimental data must be requested from the cited authors.

## Code availability

We have developed a MATLAB application named HairBundleLab to visualize and predict HB responses for any set of parameters. The application and the raw code can be accessed at our [GitHub](https://github.com/vgoyal24/HairBundleLab) repository and on MATLAB [File Exchange](https://www.mathworks.com/matlabcentral/fileexchange/178699-hairbundlelab-the-mammalian-hair-bundle-response-simulator).

