## Supplementary Information for "Three-row stereocilia model predicts mammalian hair bundle behavior"

#### This PDF file includes:

Supplementary text
Figs. S1 to S7
Tables S1 to S3
SI References

### Supplementary Information Text

### 1. Derivation of the Three-Row Micro-Mechanical Model

The dynamic equations of motion (EOM) governing the bundle's mechano-electrical response to external input form the core of this research. We start by developing a schematic of the three rows of stereocilia with necessary mechanical elements (such as springs and dampers). The schematic is illustrated in Fig. S1a and b, where a single column of three rows of stereocilia represents the multi-column HB. We designate the tallest, middle, and shortest stereocilia as row 1, row 2, and row 3, respectively. The geometric parameters are depicted in Fig. S1c, showing the bundle in an upright position. We have two adaptation complexes, or "motors", in our model that regulate fast and slow adaptation in the model.

The three degrees of freedom in our model are the bundle rotation $\phi$ (defined as the rotation of row 1), the first adaptation complex in row 1 ($l_{a1}$), and the second adaptation complex in row 2 ($l_{a2}$). In the following sections, we derive the geometric relations and the energy of each mechanical element in terms of these three generalized coordinates for two types of step inputs. The first is a direct force application (Fig. S1a), as in fluid-jet experiments (1). The second is a displacement via a stiff probe (Fig. S1b), as in (1, 2), treated as an indirect force applied to the bundle that depends on both bundle rotation and the displacement of the probe's free end, controlled experimentally by a piezo actuator.

**A. Geometric relations between rows 1 and 2.** We use the two polygons $AB'GJIBD$ and $AB'BD$ shown in Fig. S2a to derive four equations (equations 1-4) with four unknowns: $r_{B'A}$, $\alpha_2$, $l_1$, and $\alpha_3$, all expressed as functions of $\phi$ and $l_{a1}$.

$$r_{BD}\cos(\alpha_2) + (r_1 + r_2)\cos(\phi) - r_{B'A}\sin(\phi) = b_{12}, \tag{1}$$

$$(r_1 + r_2)\sin(\phi) + r_{B'A}\cos(\phi) = r_{BD}\sin(\alpha_2), \tag{2}$$

$$(r_{BD} + r_2)\sin(\alpha_2) + l_1\cos(\phi + \alpha_3) - r_1\sin(\phi) = (l_1^{gs} - l_{a1})\cos(\phi), \tag{3}$$

$$(r_{BD} + r_2)\cos(\alpha_2) + l_1\sin(\phi + \alpha_3) + r_1\cos(\phi) = (l_1^{gs} - l_{a1})\sin(\phi) + b_{12}, \tag{4}$$

where $r_{B'A}$ determines the representative distance from point $A$ to the contact point of the middle row to row 1 as if it were a stick model, and $l_1$ represents the length of the tip link between rows 1 and 2, including the gating spring. Given our model's assumption that the tip link is inextensible, any change in $l_1$ results entirely from the extension or compression of the gating spring. The system of equations in equations 1-4 yields four unknowns in terms of $\phi$ and $l_{a1}$ shown in equations 5-8.

$$r_{B'A} = -b_{12}\sin(\phi) + \sqrt{(-b_{12}\sin(\phi))^2 - [b_{12}^2 + (r_1 + r_2)^2 - 2b_{12}(r_1 + r_2)\cos(\phi) - r_{BD}^2]}, \tag{5}$$

$$\alpha_2 = \cos^{-1}\left(\frac{b_{12} + r_{B'A}\sin(\phi) - (r_1 + r_2)\cos(\phi)}{r_{BD}}\right), \tag{6}$$

$$\begin{aligned} l_1 = \{&(l_1^{gs} - l_{a1})^2 + (r_{BD} + r_2)^2 + r_1^2 + b_{12}^2 - 2(r_{BD} + r_2)(l_1^{gs} - l_{a1})\sin(\phi + \alpha_2) + 2r_1(r_{BD} + r_2)\cos(\phi + \alpha_2) \\ &- 2b_{12}\left[(r_{BD} + r_2)\cos(\alpha_2) + r_1\cos(\phi) - (l_1^{gs} - l_{a1})\sin(\phi)\right]\}^{1/2}, \end{aligned} \tag{7}$$

$$\alpha_3 = \cos^{-1}\left(\frac{(r_{BD} + r_2)\cos(\alpha_2) + r_1\cos(\phi) - b_{12} - (l_1^{gs} - l_{a1})\sin(\phi)}{l_1}\right) - \alpha_1 \tag{8}$$

**B. Geometric relations between rows 2 and 3.** From the two polygons shared between rows 2 and 3 ($DE'G'J'I'EF$ and $DE'EF$ shown in Fig. S2a), we derive four equations (equations 9-12) with four unknowns: $r_{E'D}$, $\alpha_5$, $l_2$, and $\alpha_6$ in terms of $\alpha_4$ and $l_{a2}$.

$$r_{EF}\cos(\alpha_5) + (r_2 + r_3)\sin(\alpha_4) + r_{E'D}\cos(\alpha_4) = b_{23}, \tag{9}$$

$$-(r_2 + r_3)\cos(\alpha_4) + r_{E'D}\sin(\alpha_4) = r_{EF}\sin(\alpha_5), \tag{10}$$

$$(r_{EF} + r_3)\sin(\alpha_5) + l_2\sin(\alpha_4 + \alpha_6) + r_2\cos(\alpha_4) = (l_2^{gs} - l_{a2})\sin(\alpha_4), \tag{11}$$

$$(r_{EF} + r_3)\cos(\alpha_5) - l_2\cos(\alpha_4 + \alpha_6) + r_2\sin(\alpha_4) = -(l_2^{gs} - l_{a2})\cos(\alpha_4) + b_{23}, \tag{12}$$

where $\alpha_4$ is indirectly related to the first generalized coordinate $\phi$ as $\alpha_4 = \pi - \alpha_2(\phi)$ and $\alpha_2$ is defined in equation 6. Akin to the taller stereocilia pair, $r_{E'D}$ denotes the representative height of the contact point of the shortest row to the middle row as measured from point $D$ in Fig. S2a while visualizing the system as a stick model, and $l_2$ represents the length of the tip link connecting rows 2 and 3, which varies as the corresponding gating spring extends or compresses. Solving the system of equations (equations 9-12) yields expressions for the unknowns as shown in equations 13-16.

$$r_{E'D} = b_{23}\cos(\alpha_4) + \sqrt{(b_{23}\cos(\alpha_4))^2 - [b_{23}^2 + (r_2+r_3)^2 - 2b_{23}(r_2+r_3)\sin(\alpha_4) - r_{EF}^2]}, \quad [13]$$

$$\alpha_5 = \cos^{-1}\left(\frac{b_{23} - r_{E'D}\cos(\alpha_4) - (r_2+r_3)\sin(\alpha_4)}{r_{EF}}\right), \quad [14]$$

$$l_2 = \{(l_2^{gs} - l_{a2})^2 + (r_{EF}+r_3)^2 + r_2^2 + b_{23}^2 + 2(r_{EF}+r_3)(l_2^{gs} - l_{a2})\cos(\alpha_4+\alpha_5) + 2r_2(r_{EF}+r_3)\sin(\alpha_4+\alpha_5) - 2b_{23}\left[(r_{EF}+r_3)\cos(\alpha_5) + r_2\sin(\alpha_4) + (l_2^{gs} - l_{a2})\cos(\alpha_4)\right]\}^{1/2}, \quad [15]$$

$$\alpha_6 = \cos^{-1}\left(\frac{(r_{EF}+r_3)\cos(\alpha_5) + r_2\sin(\alpha_4) - b_{23} + (l_2^{gs} - l_{a2})\cos(\alpha_4)}{l_2}\right) - \alpha_4 \quad [16]$$

**C. Defining the necessary first derivatives.** The first partial derivatives of the eight time-dependent variables computed in Sections 2A and 2B are essential for the computation of the EOM using Lagrange's methods (detailed in Section 2G). Consequently, we provide these derivatives in a tabular format in this section. Note that the complete expressions are too extensive to be written here in full and are computed using symbolic representations in MATLAB. Table S2 summarizes the derivatives and their corresponding symbols.

**D. The gating mechanism.** The gating mechanism is illustrated in Fig. S3. As the gating spring undergoes extension, it pulls the gate open at the LTLD, thereby increasing the open probability. The extent to which a gate can open is limited by a gating swing ($d_1$ or $d_2$) along each tip link (illustrated on the right in Fig. S3a). Fettiplace *et al.* (3) estimated approximately 3.5 transduction channels per tip link in apical mice. Consequently, in our model, we introduce the flexibility for the LTLD to be equipped with $N_c$ channels per tip link as illustrated for $N_c = 4$ in Fig. S3b. The open probability of a gate at each LTLD ($P_{o1}$ or $P_{o2}$) is defined by a two-state Boltzmann relation, derived by computing the difference in energies between the open and closed MET channel states, as shown in equation 17.

$$P_{oj} = \frac{1}{1 + \exp\left[-\frac{K_{GS_j} d_j}{N k_B T}(\delta l_j - l_j^{act})\right]}, \quad [17]$$

where $j = 1$ and $j = 2$ correspond to the LTLD of rows 2 and 3, respectively. $\delta l_1 = l_1 - l_1^o$ and $\delta l_2 = l_2 - l_2^o$ represent the change in length of the first and second gating springs, with $l_1$ and $l_2$ defined in equation 7 and equation 15, respectively, and $l_1^o$ and $l_2^o$ denoting the natural spring lengths defined when $\phi = l_{a1} = l_{a2} = 0$. The remaining parameters are defined in Table S1. The probability expressions in equation 17 are probability density functions, as they estimate the channel open probability of all columns in this model. As shown in Fig. S3b, each stereocilium in rows 1 and 2 has $N_c$ channel gating springs ($k_{gs_j}$) in parallel, collectively forming the stiffness of the gating spring for each channel in one column (say $\kappa_{GS_j,single} = N_c k_{gs_j}$). Due to the homogenization of the multi-column HB into a single column, $K_{GS_j} = 30\kappa_{GS_j,single}$ represents a combined value for all columns in the model, where 30 is the total number of columns in our model.

**E. Spring energies and work done by external input.** In this section, we obtain energies associated with the extension or compression of various springs in the model. The potential energies of the two extent springs and three torsional pivot springs, defined in Table S1 and Fig. S1a or b, are given by the following expressions: $U_{ES_1} = 0.5K_{ES_1}\delta l_{a1}^2$, $U_{ES_2} = 0.5K_{ES_2}\delta l_{a2}^2$, $U_{SP_1} = 0.5K_{SP_1}\delta\alpha_1^2$, $U_{SP_2} = 0.5K_{SP_2}\delta\alpha_2^2$, and $U_{SP_3} = 0.5K_{SP_3}\delta\alpha_5^2$. In all expressions, $\delta$ represents a change from the corresponding value when $\phi = l_{a1} = l_{a2} = 0$ and is represented with a superscript "$o$" (for instance, $\delta l_{a1} = l_{a1} - l_{a1}^o$). The potential function for the gating springs could not be derived in closed-form since the corresponding gating forces, $F_{GS_1} = K_{GS_1}(\delta l_1 - d_1 P_{o1})$ and $F_{GS_2} = K_{GS_2}(\delta l_2 - d_2 P_{o2})$, are not integrable due to the constitutive and geometric nonlinearities. Therefore, we treat these forces as miscellaneous force contributions in Lagrange's equations for the derivation of the EOM (refer Section 2F).

In the case of the forced HB, we define the step-force as shown in equation 18.

$$F_{ext}(t) = F_o(1 - \exp(-t/\tau_F)), \quad [18]$$

where the subscript "$F$" denotes force, $\tau_F$ represents the rise time, and $F_o$ is the amplitude of the force. Similarly, the displacement of the probe's base, $X_{ext}^{base}(t)$, features a rise time $\tau_F$ and amplitude $X_o^{base}$, as indicated in equation 19.

$$X_{ext}^{base}(t) = X_o^{base}(1 - \exp(-t/\tau_F)) \quad [19]$$

For the probe displacement, we retain the subscript "$F$" for the rise-time because it can be represented as a force shown in equation 56.

The work done by the external force ($F_{ext}$ that always acts along $x$-axis as shown in Fig. S1a at the tip of row 1) is obtained by taking a dot product of this force and the bundle rotation resolved along the Cartesian coordinates as shown in equation 20.

$$W^{ext} = -F_{ext} r_{CA} \left( \sin(\phi) - \sin(\phi^e) \right), \tag{20}$$

where $\phi^e$ denotes the resting state of the bundle, which may or may not be equal to $\phi^o$. We can obtain $\phi^e \neq \phi^o$ by applying a static load that biases the bundle to $\phi^e$.

When the HB is stimulated by a probe, as shown in Fig. S1b, we compute the potential energy stored in the spring $K_P$, representing the probe stiffness, as shown in equation 21. This potential energy depends on the relative displacement between the bundle tip and the probe's base (controlled externally by a piezo-actuator). Note that we still denote this energy by the same quantity $W^{ext}$ for generality.

$$W^{ext} = \frac{1}{2} K_P (X_{hb} - X_{ext}^{base})^2, \tag{21}$$

where $X_{hb}$ is defined as the bundle motion along the $x$-axis, a function of $\phi$, as shown in equation 42.

**F. Dissipation energy, and miscellaneous moments and forces.** In this analysis, we define the dissipation energy attributable to the motion of the bundle within the surrounding viscous fluid by presuming specific values for the damping coefficients ($\lambda_i \ \forall i \in [1,3]$ and $\lambda_{aj} \ \forall j \in [1,2]$). For the schematic shown in Fig. S1a or b, the Rayleigh dissipation function is given in equation 22.

$$\mathcal{F} = \frac{1}{2}\lambda_1 \delta\dot{\phi}^2 + \frac{1}{2}\lambda_2 \delta\dot{\alpha_2}^2 + \frac{1}{2}\lambda_3 \delta\dot{\alpha_5}^2 + \frac{1}{2}\lambda_{a1} \delta\dot{l_{a1}}^2 + \frac{1}{2}\lambda_{a2} \delta\dot{l_{a2}}^2, \tag{22}$$

where a dot $(\cdot)$ over the variables denotes the first time-derivative, geometric variables are illustrated in Fig. S1c, and the remaining parameters are listed in Table S1. The emergence of additional forces is attributed to the gating force acting along the longitudinal axes of the two gating springs. Although conservative, these forces are added separately due to their nonlinear and non-integrable nature, which precludes the derivation of a potential function. Since the first generalized coordinate is an angular dimension ($\phi$), the first EOM is a moment-balance equation. We derive the moment of $F_{GS_1}$ by using $\vec{r} \times \vec{F}$ and obtain the expression shown in equation 23.

$$M_{12} = -K_{GS_1}(\delta l_1 - d_1 P_{o1}) l_1^m, \tag{23}$$

where $l_1^m = [(l_1^{gs} - (l_{a1} - l_{a1}^o)) \sin(\alpha_3) + r_1 \cos(\alpha_3)]$ is the moment-arm length measured from the pivot point $A$, as shown in Fig. S2b, and model parameters can be found in Table S1. Similarly, the moment of the gating spring force for the spring between rows 2 and 3 ($F_{GS_2}$) is given in equation 24.

$$M_{23} = -K_{GS_2}(\delta l_2 - d_2 P_{o2}) l_2^m, \tag{24}$$

where $l_2^m = [(l_2^{gs} - (l_{a2} - l_{a2}^o)) \sin(\alpha_6) + r_2 \cos(\alpha_6) + b_{12} \cos(\alpha_2 - \alpha_6)]$ is the moment-arm length measured from the pivot point $A$ for $F_{GS_2}$, as shown in Fig. S2b with model parameters tabulated in Table S1. The second and the third EOM are force-balance equations as the remaining variables estimate length changes ($l_{a1}$ and $l_{a2}$). The component of $F_{GS_1}$ that activates the first adaptation motor is given by $Q_{12}$ in equation 25.

$$Q_{12} = -K_{GS_1}(\delta l_1 - d_1 P_{o1}) \cos(\alpha_3) \tag{25}$$

Similarly, for the gating spring between rows 2 and 3, the component of $F_{GS_2}$ that stimulates the second adaptation motor is given by $Q_{23}$ in equation 26.

$$Q_{23} = -K_{GS_2}(\delta l_2 - d_2 P_{o2}) \cos(\alpha_6) \tag{26}$$

The second set of additional forces arises from the feedback of the infiltrating cations on the adaptation motors and the stall force of the adaptation motors. Since the row 1 adaptation complex is far from either MET channel, we assume that the stall force of the first adaptation motor is not affected by the cations as given in equation 27 by $Q_{12}^{ion}$.

$$Q_{12}^{ion} = -F_{max_1}, \tag{27}$$

where $F_{max_1}$ is defined in Table S1. For the second adaptation motor site, due to its proximity to row 2 MET channels, we include an active feedback component from the infiltrating cations. The cation-mediated stall force is derived following (4) and is denoted by $Q_{23}^{ion}$ in equation 28.

$$Q_{23}^{ion} = -F_{max_2}(1 - SP_{o1}), \tag{28}$$

where parameter definitions and their values are listed in Table S1.

**G. Lagrange formulation for EOM.** Implementing Lagrange's energy method (5) is the final step in deriving the EOM for a mammalian HB. Since we have three generalized coordinates ($\phi$, $l_{a1}$, and $l_{a2}$), we employ Lagrange's energy method for each coordinate separately, as shown in equations 29-31.

$$-\frac{\partial \mathcal{F}}{\partial \dot{\phi}} + M_{12} + M_{23} = -\frac{d}{dt}\left[\frac{\partial W^{ext}}{\partial \dot{\phi}} + \sum_{i=1}^{3}\frac{U_{SP_i}}{\partial \dot{\phi}} + \sum_{j=1}^{2}\frac{U_{ES_j}}{\partial \dot{\phi}}\right] + \frac{\partial W^{ext}}{\partial \phi} + \sum_{i=1}^{3}\frac{\partial U_{SP_i}}{\partial \phi} + \sum_{j=1}^{2}\frac{\partial U_{ES_j}}{\partial \phi}, \tag{29}$$

$$-\frac{\partial \mathcal{F}}{\partial \dot{l}_{a1}} + Q_{12} + Q_{12}^{ion} = -\frac{d}{dt}\left[\frac{\partial W^{ext}}{\partial \dot{l}_{a1}} + \sum_{i=1}^{3}\frac{U_{SP_i}}{\partial \dot{l}_{a1}} + \sum_{j=1}^{2}\frac{U_{ES_j}}{\partial \dot{l}_{a1}}\right] + \frac{\partial W^{ext}}{\partial l_{a1}} + \sum_{i=1}^{3}\frac{\partial U_{SP_i}}{\partial l_{a1}} + \sum_{j=1}^{2}\frac{\partial U_{ES_j}}{\partial l_{a1}}, \tag{30}$$

$$-\frac{\partial \mathcal{F}}{\partial \dot{l}_{a2}} + Q_{23} + Q_{23}^{ion} = -\frac{d}{dt}\left[\frac{\partial W^{ext}}{\partial \dot{l}_{a2}} + \sum_{i=1}^{3}\frac{U_{SP_i}}{\partial \dot{l}_{a2}} + \sum_{j=1}^{2}\frac{U_{ES_j}}{\partial \dot{l}_{a2}}\right] + \frac{\partial W^{ext}}{\partial l_{a2}} + \sum_{i=1}^{3}\frac{\partial U_{SP_i}}{\partial l_{a2}} + \sum_{j=1}^{2}\frac{\partial U_{ES_j}}{\partial l_{a2}} \tag{31}$$

***G.1. Force input.*** Simplifying equation 29-31 using $W^{ext}$ defined in equation 20, we get

$$-\left(\lambda_1 + \lambda_2\epsilon_1^2 + \lambda_3\epsilon_7^2\right)\frac{d\phi}{dt} = K_{SP_1}(\delta\phi) + K_{SP_2}(\delta\alpha_2)\epsilon_1 + K_{SP_3}(\delta\alpha_5)\epsilon_7 - F_{ext}(t)r_{CA}\cos(\phi) - M_{12} - M_{23}, \tag{32}$$

$$-\lambda_{a1}\frac{dl_{a1}}{dt} = K_{ES_1}(\delta l_{a1}) - Q_{12} - Q_{12}^{ion}, \tag{33}$$

$$-\lambda_{a2}\frac{dl_{a2}}{dt} = K_{ES_2}(\delta l_{a2}) - Q_{23} - Q_{23}^{ion} \tag{34}$$

We denote $\lambda_1 + \lambda_2\epsilon_1^2 + \lambda_3\epsilon_7^2 = \Lambda$, a function of $\phi$, as the equivalent extracellular damping coefficient. Substituting equations 23-28 in the above equations, we obtain the final form of EOM as shown in equations 35-37.

$$\begin{aligned}\Lambda\frac{d\phi}{dt} &= -K_{GS_1}(l_1 - l_1^o - d_1P_{o1})l_1^m - K_{GS_2}(l_2 - l_2^o - d_2P_{o2})l_2^m \\ &\quad - K_{SP_1}(\phi - \phi^o) - K_{SP_2}(\alpha_2 - \alpha_2^o)\epsilon_1 - K_{SP_3}(\alpha_5 - \alpha_5^o)\epsilon_7 + F_{ext}(t)r_{CA}\cos(\phi),\end{aligned} \tag{35}$$

$$\lambda_{a1}\frac{dl_{a1}}{dt} = -K_{ES_1}(l_{a1} - l_{a1}^o) + K_{GS_1}(l_1 - l_1^o - d_1P_{o1})\cos(\alpha_3) - F_{max_1}, \tag{36}$$

$$\lambda_{a2}\frac{dl_{a2}}{dt} = -K_{ES_2}(l_{a2} - l_{a2}^o) + K_{GS_2}(l_2 - l_2^o - d_2P_{o2})\cos(\alpha_6) - F_{max_2}(1 - SP_{o1}) \tag{37}$$

***G.2. Stiff probe displacement input.*** For the displacement input model, equation 32 is altered by using $W^{ext}$ defined in equation 21. This modification yields the expression shown in equation 38 for the first equation of motion, while the remaining two, equations 36 and 37, remain unaltered.

$$-\Lambda\frac{d\phi}{dt} = K_{SP_1}(\delta\phi) + K_{SP_2}(\delta\alpha_2)\epsilon_1 + K_{SP_3}(\delta\alpha_5)\epsilon_7 + K_P r_{CA}^2\cos(\phi)\sin(\phi) - K_P X_{ext}^{base}(t)r_{CA}\cos(\phi) - M_{12} - M_{23}, \tag{38}$$

Substituting equations 23-24 in the above equation, we obtain the final form of EOM for displacement input as shown in equations 39-41.

$$\begin{aligned}\Lambda\frac{d\phi}{dt} &= -K_{GS_1}(l_1 - l_1^o - d_1P_{o1})l_1^m - K_{GS_2}(l_2 - l_2^o - d_2P_{o2})l_2^m \\ &\quad - K_{SP_1}(\phi - \phi^o) - K_{SP_2}(\alpha_2 - \alpha_2^o)\epsilon_1 - K_{SP_3}(\alpha_5 - \alpha_5^o)\epsilon_7 - K_P r_{CA}^2\cos(\phi)\sin(\phi) + K_P X_{ext}^{base}(t)r_{CA}\cos(\phi),\end{aligned} \tag{39}$$

$$\lambda_{a1}\frac{dl_{a1}}{dt} = -K_{ES_1}(l_{a1} - l_{a1}^o) + K_{GS_1}(l_1 - l_1^o - d_1P_{o1})\cos(\alpha_3) - F_{max_1}, \tag{40}$$

$$\lambda_{a2}\frac{dl_{a2}}{dt} = -K_{ES_2}(l_{a2} - l_{a2}^o) + K_{GS_2}(l_2 - l_2^o - d_2P_{o2})\cos(\alpha_6) - F_{max_2}(1 - SP_{o1}) \tag{41}$$

***G.3. Model outputs.*** The motion of the bundle along the $x$-axis or the mechanical response of the HB is given by

$$X_{hb} = r_{CA}\sin(\phi), \quad [42]$$

and the MET current or the electrical response of the HB is given by

$$I_{met} = \frac{G_{max}(dV_{hb}^o - EP)}{1+\exp\left[-\frac{K_{GS_1}d_1}{Nk_BT}(\delta l_1 - l_1^{act})\right]} + \frac{G_{max}(dV_{hb}^o - EP)}{1+\exp\left[-\frac{K_{GS_2}d_2}{Nk_BT}(\delta l_2 - l_2^{act})\right]}, \quad [43]$$

where the mechanical and electrical parameter definitions and values are tabulated in Table S1.

**H. Bundle stiffness.** The stiffness of the bundle is computed from both fluid-jet and stiff probe models under two conditions. The first condition is when the tip links are intact, in which case we use the complete nonlinear model as described in equations 35-37 for the fluid-jet model and equations 39-41 for the stiff probe model. The second condition is when the tip links are disrupted, which results in the inactivation of channel transduction and the two adaptive mechanisms. This second condition emulates the *in-vivo* HB when the introduction of different external compounds like BAPTA (6) or EDTA (7) severs the tip links.

***H.1. Intact tip links force input model.*** To derive the bundle stiffness analytically from this model, we use equation 35. We can represent the external force at the steady-state ($d(\cdot)/dt = 0$) as shown in equation 44.

$$F_{ext} = \frac{1}{r_{CA}\cos(\phi)}[K_{GS_1}(l_1 - l_1^o - d_1P_{o1})l_1^m + K_{GS_2}(l_2 - l_2^o - d_2P_{o2})l_2^m + K_{SP_1}(\phi - \phi^o) + K_{SP_2}(\alpha_2 - \alpha_2^o)\epsilon_1 + K_{SP_3}(\alpha_5 - \alpha_5^o)\epsilon_7] \quad [44]$$

The bundle stiffness is given by $K_{HB} = dF_{ext}/dX_{hb}$, where $X_{hb} = r_{CA}\sin(\phi)$. Since $F_{ext}$ depends on all three degrees of freedom, we can write the expression for bundle stiffness in the form of a total derivative, as shown in equation 45 following the chain rule of differentiation.

$$\frac{dF_{ext}}{dX_{hb}} = \frac{\partial\phi}{\partial X_{hb}}\frac{\partial F_{ext}}{\partial\phi} + \frac{\partial l_{a1}}{\partial X_{hb}}\frac{\partial F_{ext}}{\partial l_{a1}} + \frac{\partial l_{a2}}{\partial X_{hb}}\frac{\partial F_{ext}}{\partial l_{a2}} = \frac{\partial\phi}{\partial X_{hb}}\frac{\partial F_{ext}}{\partial\phi} + \frac{\partial\phi}{\partial X_{hb}}\frac{\partial l_{a1}}{\partial\phi}\frac{\partial F_{ext}}{\partial l_{a1}} + \frac{\partial\phi}{\partial X_{hb}}\frac{\partial l_{a2}}{\partial\phi}\frac{\partial F_{ext}}{\partial l_{a2}} \quad [45]$$

Note that $\partial\phi/\partial X_{hb} = 1/r_{CA}\cos(\phi)$ using equation 42. Equation 46 gives the complete analytical expression for the bundle stiffness.

$$K_{HB} = \frac{1}{r_{CA}\cos(\phi)}\left[\frac{\partial F_{ext}}{\partial\phi} + \frac{\partial l_{a1}}{\partial\phi}\frac{\partial F_{ext}}{\partial l_{a1}} + \frac{\partial l_{a2}}{\partial\phi}\frac{\partial F_{ext}}{\partial l_{a2}}\right], \quad [46]$$

where the RHS must be evaluated at $(\phi^{ss}, l_{a1}^{ss}, l_{a2}^{ss})$ obtained from nonlinear simulations at the steady-state "$ss$" corresponding to each applied force or displacement amplitude. The final step is to find $\partial l_{a1}/\partial\phi$ and $\partial l_{a2}/\partial\phi$. For that, we use equations 36 and 37. The derivative of equation 36 and equation 37 with respect to $\phi$ at the steady-state is defined in equation 47 and equation 48, respectively.

$$0 = \frac{d}{d\phi}\left[-K_{ES_1}(l_{a1} - l_{a1}^o) + K_{GS_1}(l_1 - l_1^o - d_1P_{o1})\cos(\alpha_3) - F_{max_1}\right] \quad [47]$$

$$0 = \frac{d}{d\phi}\left[-K_{ES_2}(l_{a2} - l_{a2}^o) + K_{GS_2}(l_2 - l_2^o - d_2P_{o2})\cos(\alpha_6) - F_{max_2}(1 - SP_{o1})\right] \quad [48]$$

Let us denote the expression to be differentiated in equation 47 as $l_{a1}^{RHS}$ and that in equation 48 as $l_{a2}^{RHS}$, so that equations 47 and 48 modify to equations 49 and 50, respectively. Further, $l_{a1}^{RHS}$ and $l_{a2}^{RHS}$ depend on all three degrees of freedom. Therefore, we apply the chain rule as shown in the same equations.

$$0 = \frac{dl_{a1}^{RHS}}{d\phi} = \frac{\partial l_{a1}^{RHS}}{\partial\phi} + \frac{\partial l_{a1}}{\partial\phi}\frac{\partial l_{a1}^{RHS}}{\partial l_{a1}} + \frac{\partial l_{a2}}{\partial\phi}\frac{\partial l_{a1}^{RHS}}{\partial l_{a2}} \quad [49]$$

$$0 = \frac{dl_{a2}^{RHS}}{d\phi} = \frac{\partial l_{a2}^{RHS}}{\partial\phi} + \frac{\partial l_{a1}}{\partial\phi}\frac{\partial l_{a2}^{RHS}}{\partial l_{a1}} + \frac{\partial l_{a2}}{\partial\phi}\frac{\partial l_{a2}^{RHS}}{\partial l_{a2}} \quad [50]$$

Now, we can generate a matrix-vector system to compute $\partial l_{a1}/\partial\phi$ and $\partial l_{a2}/\partial\phi$ as shown in equation 51.

$$\begin{bmatrix} \frac{\partial l_{a1}^{RHS}}{\partial l_{a1}} & \frac{\partial l_{a1}^{RHS}}{\partial l_{a2}} \\ \frac{\partial l_{a2}^{RHS}}{\partial l_{a1}} & \frac{\partial l_{a2}^{RHS}}{\partial l_{a2}} \end{bmatrix} \begin{bmatrix} \frac{\partial l_{a1}}{\partial\phi} \\ \frac{\partial l_{a2}}{\partial\phi} \end{bmatrix} = \begin{bmatrix} -\frac{\partial l_{a1}^{RHS}}{\partial\phi} \\ -\frac{\partial l_{a2}^{RHS}}{\partial\phi} \end{bmatrix} \quad [51]$$

Finally, we take an inverse of the $2 \times 2$ matrix to obtain functional expressions of $\partial l_{a1}/\partial\phi$ and $\partial l_{a2}/\partial\phi$ as shown in equation 52.

$$\begin{bmatrix} \dfrac{\partial l_{a1}}{\partial\phi} \\ \dfrac{\partial l_{a2}}{\partial\phi} \end{bmatrix} = \begin{bmatrix} \dfrac{\partial l_{a1}^{RHS}}{\partial l_{a1}} & \dfrac{\partial l_{a1}^{RHS}}{\partial l_{a2}} \\ \dfrac{\partial l_{a2}^{RHS}}{\partial l_{a1}} & \dfrac{\partial l_{a2}^{RHS}}{\partial l_{a2}} \end{bmatrix}^{-1} \begin{bmatrix} -\dfrac{\partial l_{a1}^{RHS}}{\partial\phi} \\ -\dfrac{\partial l_{a2}^{RHS}}{\partial\phi} \end{bmatrix} \qquad [52]$$

Equation 52 can be substituted in equation 46 to obtain an analytical expression for $K_{HB}$ in terms of the three degrees of freedom and the model parameters. Note that the partial derivatives of $F_{ext}$, $l_{a1}^{RHS}$, and $l_{a2}^{RHS}$ are complex symbolic expressions computed using MATLAB's symbolic toolbox, and are therefore not explicitly written here. But, they can be derived by hand calculations.

To compute the stiffness numerically from the time-domain nonlinear solutions, we obtain the steady-state HB displacements (nearing the end of the force stimulus) corresponding to each externally applied force amplitude $F_o$ in equation 18. The numerical bundle stiffness $K_{HB}$ is given in equation 53.

$$K_{HB} = \frac{F_o^{k+1} - F_o^k}{X_{hb}^{k+1} - X_{hb}^k}, \qquad [53]$$

where $k$ represents the $k^{th}$ force level.

***H.2. Disrupted tip links force input model.*** In this model, there are no active transduction channels due to the disruption of the tip links. As a result, there are no gating and extent springs in the model. Only one equation of motion corresponding to $\phi$ governs the bundle's response. Equation 44 is modified to equation 54.

$$F_{ext} = \frac{1}{r_{CA}\cos(\phi)}[K_{SP_1}(\phi - \phi^o) + K_{SP_2}(\alpha_2 - \alpha_2^o)\epsilon_1 + K_{SP_3}(\alpha_5 - \alpha_5^o)\epsilon_7] \qquad [54]$$

In this case, the analytical expression for $K_{HB}$ can be directly calculated as shown in equation 55, and the numerical value for the bundle stiffness can be obtained in the same way as shown in equation 53.

$$K_{HB} = \frac{1}{r_{CA}\cos(\phi)}\frac{dF_{ext}}{d\phi} \qquad [55]$$

***H.3. Intact tip links stiff probe displacement input model.*** For this model, we use equations 39-41. The force that the probe applies at the tip of the tallest row is differently written than that in the fluid-jet model and is given in equation 56.

$$F_{ext} = K_P(X_{ext}^{base} - X_{hb}) \qquad [56]$$

We manipulate equation 39 at the steady-state so that it incorporates equation 56, as shown in equation 57.

$$\begin{aligned} 0 = &-K_{GS_1}(l_1 - l_1^o - d_1P_{o1})l_1^m - K_{GS_2}(l_2 - l_2^o - d_2P_{o2})l_2^m \\ &- K_{SP_1}(\phi - \phi^o) - K_{SP_2}(\alpha_2 - \alpha_2^o)\epsilon_1 - K_{SP_3}(\alpha_5 - \alpha_5^o)\epsilon_7 + K_P\left[X_{ext}^{base} - r_{CA}\sin(\phi)\right] r_{CA}\cos(\phi) \end{aligned} \qquad [57]$$

Substituting equation 56 in equation 57 shows that this model is the same as the fluid-jet model for stiffness calculation. It makes sense because the probe stiffness is an external element that should not affect the inherent stiffness of the bundle. Therefore, we follow the same procedure as in Section H.1 to obtain the analytical expression for $K_{HB}$ shown in equation 46.

To compute $K_{HB}$ numerically, we use equation 58.

$$K_{HB} = K_P\frac{(X_o^{base,k+1} - X_{hb}^{k+1}) - (X_o^{base,k} - X_{hb}^k)}{X_{hb}^{k+1} - X_{hb}^k} = K_P\frac{X_o^{base,k+1} - X_o^{base,k}}{X_{hb}^{k+1} - X_{hb}^k} - K_P, \qquad [58]$$

where $X_o$ is the probe displacement amplitude as shown in equation 19 and $k$ represents its $k^{th}$ value.

***H.4. Disrupted tip links stiff probe displacement input model.*** In this final model for evaluating bundle stiffness, there are no gating or extent springs due to the absence of the tip links. The analytical expression for $K_{HB}$ is the same as in equation 55 with $F_{ext}$ from equation 56, and the analytical expression follows from equation 58.

**I. Geometric gain.** The geometric gain is defined as the ratio of the change in length of the gating spring to the displacement of the row 1 tip. From our model, since we have two gating springs from three rows, we obtain two geometric gains, labeled as $\gamma_1$ for the gating spring between rows 1 and 2 and $\gamma_2$ for the gating spring between rows 2 and 3. The two geometric gains are mathematically defined in equation 59 and equation 60, respectively.

$$\gamma_1 = \frac{\partial(\delta l_1)}{\partial X_{hb}} = \frac{\epsilon_4}{r_{CA}\cos(\phi)}, \qquad [59]$$

$$\gamma_2 = \frac{\partial(\delta l_2)}{\partial X_{hb}} = \frac{\epsilon_{10}}{r_{CA}\cos(\phi)}, \qquad [60]$$

where $\epsilon_4$ and $\epsilon_{10}$ are defined in Table S2, and $r_{CA}$ is the length of row 1 as illustrated in Fig. S1c and Table S1.

### 2. Linearized Three-Row Model

The fully nonlinear three-row model simulates responses effectively across various stimuli but obscures various underlying mechanisms that can explain experimental observations. Linearizing the model helps bridge that gap. To linearize our three-row model described in Section 1, first, we apply a static biasing load ($F_{st}$) or probe displacement ($X_{st}^{base}$) on the model to achieve a desired average resting open probability ($\sim 7\%$ here to match experimental data from (1)), providing estimates for the equilibrium values of the three coordinates, $\phi^e$, $l_{a1}^e$, and $l_{a2}^e$.

**A. Linearizing the force input model.** After achieving the desired resting open probability, we represent equations 35-37 in simpler function notations as follows:

$$\frac{d\phi}{dt} = f_1(\phi, l_{a1}) + f_2(\phi, l_{a2}) + K(\phi) + F_{ext}(t) f_3(\phi), \quad [61]$$

$$\frac{dl_{a1}}{dt} = g_1(\phi, l_{a1}) + g_2(l_{a1}) + g_3, \quad [62]$$

$$\frac{dl_{a2}}{dt} = m_1(\phi, l_{a2}) + m_2(l_{a2}) + m_3(\phi, l_{a1}), \quad [63]$$

where

$$f_1(\phi, l_{a1}) = -\frac{K_{GS_1}}{\Lambda}(l_1 - l_1^o - d_1 P_{o1}) l_1^m, \quad [64]$$

$$f_2(\phi, l_{a2}) = -\frac{K_{GS_2}}{\Lambda}(l_2 - l_2^o - d_2 P_{o2}) l_2^m, \quad [65]$$

$$K(\phi) = -\frac{1}{\Lambda}\left[K_{SP_1}(\phi - \phi^o) + K_{SP_2}(\alpha_2 - \alpha_2^o)\epsilon_1 + K_{SP_3}(\alpha_5 - \alpha_5^o)\epsilon_7\right], \quad [66]$$

$$f_3(\phi) = \frac{r_{CA}\cos(\phi)}{\Lambda}, \quad [67]$$

$$g_1(\phi, l_{a1}) = \frac{K_{GS_1}}{\lambda_{a1}}(l_1 - l_1^o - d_1 P_{o1})\cos(\alpha_3), \quad [68]$$

$$g_2(l_{a1}) = -\frac{K_{ES_1}}{\lambda_{a1}}(l_{a1} - l_{a1}^o), \quad [69]$$

$$g_3 = -\frac{F_{max_1}}{\lambda_{a1}}, \quad [70]$$

$$m_1(\phi, l_{a2}) = \frac{K_{GS_2}}{\lambda_{a2}}(l_2 - l_2^o - d_2 P_{o2})\cos(\alpha_6), \quad [71]$$

$$m_2(l_{a2}) = -\frac{K_{ES_2}}{\lambda_{a2}}(l_{a2} - l_{a2}^o), \quad [72]$$

$$m_3(\phi, l_{a1}) = -\frac{F_{max_2}}{\lambda_{a2}}(1 - SP_{o1}), \quad [73]$$

Using first-order Taylor's expansion about the equilibrium ($\phi^e$, $l_{a1}^e$, and $l_{a2}^e$) corresponding to a static, biasing load $F_{st}$, for equation 61, we get,

$$\frac{d}{dt}(\phi^e + \Delta\phi) = f_1(\phi^e, l_{a1}^e) + \Delta f_1 + f_2(\phi^e, l_{a2}^e) + \Delta f_2 + K(\phi^e) + \Delta K + (F_{st} + \Delta F_{ext}(t))(f_3(\phi^e) + \Delta f_3) \quad [74]$$

$$\implies \frac{d}{dt}(\Delta\phi) = f_1(\phi^e, l_{a1}^e) + f_2(\phi^e, l_{a2}^e) + K(\phi^e) + F_{st} f_3(\phi^e)$$
$$+ \Delta f_1 + \Delta f_2 + \Delta K + F_{st}\Delta f_3 + \Delta F_{ext}(t) f_3(\phi^e) + \Delta F_{ext}\Delta f_3(t), \quad [75]$$

where $\Delta(\cdot)$ has a general form given by the sum of partial derivatives with respect to the generalized coordinates $(\phi,\, l_{a1},\, l_{a2})$. As an example, if we represent the three generalized coordinates $\phi$, $l_{a1}$, $l_{a2}$ by $q_1$, $q_2$, $q_3$, respectively, then the general form of $\Delta(\cdot)$ is given by

$$\Delta(\cdot) = \sum_{i=1}^{3} \left.\frac{\partial(\cdot)}{\partial q_i}\right|_e \tilde{q}_i, \qquad [76]$$

where $\tilde{q}_i$ is the change in the generalized coordinate from the equilibrium position, i.e., $\tilde{q}_i = q_i - q_i^e$ (for instance, for $q_1 = \phi$, $\tilde{\phi} = \phi - \phi^e$) and "$e$" denotes variable evaluated at the equilibrium. In equation 75, we neglect the last term $\Delta F_{ext}\Delta f_3(t)$ because it is a second-order term that is infinitesimally small. At equilibrium, $f_1(\phi^e, l_{a1}^e) + f_2(\phi^e, l_{a1}^e, l_{a2}^e) + K(\phi^e) + F_{st} f_3(\phi^e) = 0$, therefore, equation 75 can be simplified to

$$\frac{d}{dt}(\Delta\phi) = \Delta f_1 + \Delta f_2 + \Delta K + F_{st}\Delta f_3 + \Delta F_{ext}(t) f_3(\phi^e) \qquad [77]$$

Note that for force, $\Delta F_{ext}(t)$ is just the perturbation above the static force as a function of time. So, we substitute $\Delta F_{ext}(t) = f_{ext}(t)$. Therefore, equation 77, after expansion becomes

$$\frac{d\tilde{\phi}}{dt} = \left.\frac{\partial f_1}{\partial \phi}\right|_e \tilde{\phi} + \left.\frac{\partial f_1}{\partial l_{a1}}\right|_e \tilde{l}_{a1} + \left.\frac{\partial f_2}{\partial \phi}\right|_e \tilde{\phi} + \left.\frac{\partial f_2}{\partial l_{a2}}\right|_e \tilde{l}_{a2} + \left.\frac{\partial K}{\partial \phi}\right|_e \tilde{\phi} + F_{st}\left.\frac{\partial f_3}{\partial \phi}\right|_e \tilde{\phi} + f_{ext}(t) f_3(\phi^e) \qquad [78]$$

Next, we take the first adaptation motor equation given in equation 62 and write its first-order Taylor's expansion.

$$\frac{d}{dt}(l_{a1}^e + \Delta l_{a1}) = g_1(\phi^e, l_{a1}^e) + \Delta g_1 + g_2(l_{a1}^e) + \Delta g_2 + g_3 + \Delta g_3 \qquad [79]$$

$$\implies \frac{d}{dt}(\Delta l_{a1}) = g_1(\phi^e, l_{a1}^e) + g_2(l_{a1}^e) + g_3 + \Delta g_1 + \Delta g_2 \qquad [80]$$

Because $g_3$ is a constant, it sets $\Delta g_3 = 0$, omitted from equation 80. At equilibrium, $g_1(\phi^e, l_{a1}^e) + g_2(l_{a1}^e) + g_3 = 0$. Using equation 76, we get

$$\frac{d}{dt}(\Delta l_{a1}) = \left.\frac{\partial g_1}{\partial \phi}\right|_e \tilde{\phi} + \left.\frac{\partial g_1}{\partial l_{a1}}\right|_e \tilde{l}_{a1} + \left.\frac{\partial g_2}{\partial l_{a1}}\right|_e \tilde{l}_{a1} \qquad [81]$$

Finally, we linearize the third equation of motion governing the dynamics of the second adaptation motor shown in equation 63.

$$\frac{d}{dt}(l_{a2}^e + \Delta l_{a2}) = m_1(\phi^e, l_{a2}^e) + \Delta m_1 + m_2(l_{a2}^e) + \Delta m_2 + m_3(\phi^e, l_{a1}^e) + \Delta m_3$$

$$\implies \frac{d}{dt}(\Delta l_{a2}) = m_1(\phi^e, l_{a2}^e) + m_2(l_{a2}^e) + m_3(\phi^e, l_{a1}^e) + \Delta m_1 + \Delta m_2 + \Delta m_3 \qquad [82]$$

At equilibrium, $m_1(\phi^e, l_{a2}^e) + m_2(l_{a2}^e) + m_3(\phi^e, l_{a1}^e) = 0$. Using equation 76, we get

$$\frac{dl_{a2}}{dt} = \left.\frac{\partial m_1}{\partial \phi}\right|_e \tilde{\phi} + \left.\frac{\partial m_1}{\partial l_{a2}}\right|_e \tilde{l}_{a2} + \left.\frac{\partial m_2}{\partial l_{a2}}\right|_e \tilde{l}_{a2} + \left.\frac{\partial m_3}{\partial \phi}\right|_e \tilde{\phi} + \left.\frac{\partial m_3}{\partial l_{a1}}\right|_e \tilde{l}_{a1} \qquad [83]$$

Now, the general matrix-vector form can be written using equation 78, equation 81, and equation 83. This yields the linear system as

$$\dot{\boldsymbol{q}} = \boldsymbol{\mathcal{J}}\boldsymbol{q} + \boldsymbol{f_{ext}}(t) \qquad [84]$$

where the three vectors are

$$\boldsymbol{q} = \begin{pmatrix} \tilde{\phi} \\ \tilde{l}_{a1} \\ \tilde{l}_{a2} \end{pmatrix},\ \dot{\boldsymbol{q}} = \frac{d}{dt}\begin{pmatrix} \tilde{\phi} \\ \tilde{l}_{a1} \\ \tilde{l}_{a2} \end{pmatrix},\ \text{and } \boldsymbol{f_{ext}}(t) = f_{ext}(t) f_3(\phi^e)\begin{pmatrix} 1 \\ 0 \\ 0 \end{pmatrix} \qquad [85]$$

and the Jacobian $\boldsymbol{\mathcal{J}}$, evaluated at the equilibrium is given by

$$\boldsymbol{\mathcal{J}} = \begin{bmatrix} \frac{\partial}{\partial \phi}(f_1 + f_2 + K + F_{st} f_3) & \frac{\partial f_1}{\partial l_{a1}} & \frac{\partial f_2}{\partial l_{a2}} \\ \frac{\partial g_1}{\partial \phi} & \frac{\partial}{\partial l_{a1}}(g_1 + g_2) & 0 \\ \frac{\partial}{\partial \phi}(m_1 + m_3) & \frac{\partial m_3}{\partial l_{a1}} & \frac{\partial}{\partial l_{a2}}(m_1 + m_2) \end{bmatrix}_e \qquad [86]$$

**B. Linearizing the stiff probe displacement input model.** After applying a biasing probe displacement, we follow the same procedure as in the previous section to linearize equations 39-41. In this case, we only need to linearize equation 39 as the linearized form of the two remaining EOM are derived in equations 81 and 83. Let the simpler function notation for equation 39 be the following:

$$\frac{d\phi}{dt} = f_1(\phi, l_{a1}) + f_2(\phi, l_{a2}) + K(\phi) + K_{pr}(\phi) + K_P X_{ext}^{base}(t) f_3(\phi), \quad [87]$$

where

$$K_{pr}(\phi) = -\frac{K_P r_{CA}^2}{\Lambda} \cos(\phi)\sin(\phi), \quad [88]$$

Using first-order Taylor's expansion about the equilibrium ($\phi^e$, $l_{a1}^e$, and $l_{a2}^e$) corresponding to a static, biasing probe displacement $X_{st}^{base}$, for equation 87, we get,

$$\begin{aligned}\frac{d}{dt}(\phi^e + \Delta\phi) = f_1(\phi^e, l_{a1}^e) + \Delta f_1 + f_2(\phi^e, l_{a2}^e) + \Delta f_2 + K(\phi^e) + \Delta K \\ + K_{pr}(\phi^e) + \Delta K_{pr} + K_P(X_{st}^{base} + \Delta X_{ext}^{base}(t))(f_3(\phi^e) + \Delta f_3)\end{aligned} \quad [89]$$

$$\begin{aligned}\implies \frac{d}{dt}(\Delta\phi) = f_1(\phi^e, l_{a1}^e) + f_2(\phi^e, l_{a2}^e) + K(\phi^e) + K_{pr}(\phi^e) + K_P X_{st}^{base} f_3(\phi^e) \\ + \Delta f_1 + \Delta f_2 + \Delta K + \Delta K_{pr} + K_P X_{st}^{base} \Delta f_3 + K_P \Delta X_{ext}^{base}(t) f_3(\phi^e) + K_P \Delta X_{ext}^{base}(t) \Delta f_3(t),\end{aligned} \quad [90]$$

where $\Delta(\cdot)$ is defined in equation 76. In equation 90, we neglect the last term $K_P \Delta X_{ext}^{base}(t)\Delta f_3(t)$ because it is a second-order term and therefore infinitesimally small for $\phi \ll 1$. At equilibrium, $f_1(\phi^e, l_{a1}^e) + f_2(\phi^e, l_{a1}^e, l_{a2}^e) + K(\phi^e) + K_{pr}(\phi^e) + K_P X_{st}^{base} f_3(\phi^e) = 0$, therefore equation 90 can be simplified to

$$\frac{d}{dt}(\Delta\phi) = \Delta f_1 + \Delta f_2 + \Delta K + \Delta K_{pr} + K_P X_{st}^{base} \Delta f_3 + K_P \Delta X_{ext}^{base}(t) f_3(\phi^e) \quad [91]$$

Note that for probe displacement, $\Delta X_{ext}^{base}(t)$ is the perturbation about the static probe displacement as a function of time. So, we substitute $\Delta X_{ext}^{base}(t) = x_{ext}^{base}(t)$. Therefore, equation 91, after expansion becomes

$$\frac{d\tilde{\phi}}{dt} = \left.\frac{\partial f_1}{\partial \phi}\right|_e \tilde{\phi} + \left.\frac{\partial f_1}{\partial l_{a1}}\right|_e \tilde{l}_{a1} + \left.\frac{\partial f_2}{\partial \phi}\right|_e \tilde{\phi} + \left.\frac{\partial f_2}{\partial l_{a2}}\right|_e \tilde{l}_{a2} + \left.\frac{\partial K}{\partial \phi}\right|_e \tilde{\phi} + \left.\frac{\partial K_{pr}}{\partial \phi}\right|_e \tilde{\phi} + K_P X_{st}^{base} \left.\frac{\partial f_3}{\partial \phi}\right|_e \tilde{\phi} + K_P x_{ext}^{base}(t) f_3(\phi^e) \quad [92]$$

Now, the general matrix-vector form for this case can be written using equation 92, equation 81, and equation 83. This yields the linear system as

$$\dot{\boldsymbol{q}} = \boldsymbol{\mathcal{J}}\boldsymbol{q} + \boldsymbol{f_{ext}}(t) \quad [93]$$

where the two vectors $\boldsymbol{q}$ and $\dot{\boldsymbol{q}}$ are defined in equation 85, and

$$\boldsymbol{f_{ext}}(t) = K_P x_{ext}^{base}(t) f_3(\phi^e) \begin{pmatrix} 1 \\ 0 \\ 0 \end{pmatrix} \quad [94]$$

Note that this linearized system (equation 93) is equivalent to the one derived in the previous section with a force input in equation 84. However, the definition of $\boldsymbol{f_{ext}}(t)$ itself has changed, along with the Jacobian. The new Jacobian $\boldsymbol{\mathcal{J}}$, evaluated at the equilibrium is given by

$$\boldsymbol{\mathcal{J}} = \begin{bmatrix} \frac{\partial}{\partial \phi}(f_1 + f_2 + K + K_{pr} + K_P X_{st}^{base} f_3) & \frac{\partial f_1}{\partial l_{a1}} & \frac{\partial f_2}{\partial l_{a2}} \\ \frac{\partial g_1}{\partial \phi} & \frac{\partial}{\partial l_{a1}}(g_1 + g_2) & 0 \\ \frac{\partial}{\partial \phi}(m_1 + m_3) & \frac{\partial m_3}{\partial l_{a1}} & \frac{\partial}{\partial l_{a2}}(m_1 + m_2) \end{bmatrix}_e \quad [95]$$

From the linear system in equation 84 or equation 93, we obtain three time constants by evaluating the eigenvalues of the $3 \times 3$ matrix $\boldsymbol{\mathcal{J}}$ as functions of the model parameters at equilibrium. The solution of this linear system comprises the homogeneous solution and the particular solution corresponding to the externally applied force or displacement (general step-input shown in equation 105). In the subsequent sections, we derive the closed-form solution to the linearized system by defining the initial conditions as $\tilde{\phi}(t = 0) = \tilde{l}_{a1}(t = 0) = \tilde{l}_{a2}(t = 0) = 0$.

**C. Homogeneous solution to the linearized model.** In the process of obtaining the solution to the linearized system in equation 84 and equation 93, we begin by evaluating the homogeneous solution first, i.e., when $\boldsymbol{f_{ext}}(t) = \boldsymbol{x_{ext}^{base}}(t) = \mathbf{0}$. The homogeneous system is given by

$$\dot{\boldsymbol{q}} = \boldsymbol{\mathcal{J}}\boldsymbol{q} \tag{96}$$

Let the general solution be $\boldsymbol{q} = \boldsymbol{u}\exp(\zeta t)$, where $\boldsymbol{u}$ is an eigenvector and $\zeta$ is the corresponding eigenvalue. Substituting the general solution in equation 96, we obtain the characteristic equation shown in equation 99.

$$\zeta\boldsymbol{u}\exp(\zeta t) = \boldsymbol{\mathcal{J}}\boldsymbol{u}\exp(\zeta t) \tag{97}$$

$$\implies (\boldsymbol{\mathcal{J}} - \zeta\boldsymbol{\mathcal{I}})\boldsymbol{u} = 0 \tag{98}$$

$$\implies \det(\boldsymbol{\mathcal{J}} - \zeta\boldsymbol{\mathcal{I}}) = 0, \tag{99}$$

where $\boldsymbol{\mathcal{I}}$ is a $3 \times 3$ identity matrix. Solving the characteristic equations yields three eigenvalues ($\zeta_1$, $\zeta_2$, and $\zeta_3$) and a $3 \times 3$ eigenvector matrix given by $\mathbf{V}$ composed of the three eigenvectors ($\boldsymbol{u^1}$, $\boldsymbol{u^2}$, and $\boldsymbol{u^3}$), as shown in equation 100. Note that since the Jacobian matrix in the case of force input (equation 86) is different from the displacement input (equation 95), the eigenvalues and eigenvectors would be different.

$$\mathbf{V} = \begin{pmatrix} \vdots & \vdots & \vdots \\ \boldsymbol{u^1} & \boldsymbol{u^2} & \boldsymbol{u^3} \\ \vdots & \vdots & \vdots \end{pmatrix} \tag{100}$$

The homogeneous solution can be evaluated using equation 101.

$$\boldsymbol{q}(t) = \mathbf{V}e^{\mathbf{J}t}\mathbf{c}, \tag{101}$$

where $e^{\mathbf{J}t}$ is the matrix exponential of the third-order diagonal Jordan matrix $\mathbf{J}$ formed by the eigenvalues and $\mathbf{c} = \{c_1, c_2, c_3\}^T$ represents three constants that depend on the initial conditions. This computation gives us three equations (equations 102-104) with three constant unknowns ($c_1$, $c_2$, and $c_3$).

$$\tilde{\phi}^h = c_1 u_1^1 e^{\zeta_1 t} + c_2 u_1^2 e^{\zeta_2 t} + c_3 u_1^3 e^{\zeta_3 t}, \tag{102}$$

$$\tilde{l}_{a1}^h = c_1 u_2^1 e^{\zeta_1 t} + c_2 u_2^2 e^{\zeta_2 t} + c_3 u_2^3 e^{\zeta_3 t}, \tag{103}$$

$$\tilde{l}_{a2}^h = c_1 u_3^1 e^{\zeta_1 t} + c_2 u_3^2 e^{\zeta_2 t} + c_3 u_3^3 e^{\zeta_3 t}, \tag{104}$$

where superscript "$h$" denotes homogeneous solution.

**D. Particular solution in response to the external input.** We stimulate the HB with a step-input ($f_{ext}(t)$) that consists of a rise time ($\tau_F$) and a steady-state amplitude ($F_o$). Therefore, its general expression is

$$f_{ext}(t) = F_o(1 - e^{-t/\tau_F}) \tag{105}$$

For clarity, in case of force input (fluid-jet), $f_{ext}(t)$ is represented in equation 105 in its true form, whereas for a displacement input, $f_{ext}(t) = K_P x_{ext}^{base}(t)$ and $F_o = K_P X_o^{base}$. This input comprises two components: one is the steady-state value $F_o$, and the second is its exponentially decaying part $-F_o\exp(-t/\tau_F)$. Therefore, we treat them separately to evaluate the particular solution. For the steady-state (constant) input, let the solution be $\boldsymbol{q} = \boldsymbol{q}^{p_1}$, where all three elements of $\boldsymbol{q}^{p_1}$ are independent of time (implying $\dot{\boldsymbol{q}}^{p_1} = 0$). Substituting the solution in equation 84 (or equation 93), we get

$$\boldsymbol{q}^{p_1} = -\boldsymbol{\mathcal{J}}^{-1}\begin{pmatrix} F_o f_3(\phi^e) \\ 0 \\ 0 \end{pmatrix} \tag{106}$$

For the exponential force, let us assume that the solution is $\boldsymbol{q} = \boldsymbol{q}^{p_2}\exp(-t/\tau_F)$ with all three elements of $\boldsymbol{q}^{p_2}$ independent of time. Substituting the assumed solution in equation 84 (or equation 93), we obtain

$$-\frac{1}{\tau_F}\boldsymbol{\mathcal{I}}\boldsymbol{q}^{p_2}e^{-t/\tau_F} = \boldsymbol{\mathcal{J}}\boldsymbol{q}^{p_2}e^{-t/\tau_F} - \begin{pmatrix} F_o f_3(\phi^e) \\ 0 \\ 0 \end{pmatrix} e^{-t/\tau_F} \tag{107}$$

$$\implies \boldsymbol{q}^{p_2} = \left(\boldsymbol{\mathcal{J}} + \frac{1}{\tau_F}\boldsymbol{\mathcal{I}}\right)^{-1}\begin{pmatrix} F_o f_3(\phi^e) \\ 0 \\ 0 \end{pmatrix} \tag{108}$$

**E. Complete solution to the linearized model.** Now that we have the homogeneous and the particular solutions, we can write the complete solution by adding equations 102-104, equation 106, and equation 108 as shown in equations 109-111.

$$\tilde{\phi}(t) = c_1 u_1^1 e^{-t/\tau_1} + c_2 u_1^2 e^{-t/\tau_2} + c_3 u_1^3 e^{-t/\tau_3} + q_1^{p_1} + q_1^{p_2} e^{-t/\tau_F}, \quad [109]$$

$$\tilde{l}_{a1}(t) = c_1 u_2^1 e^{-t/\tau_1} + c_2 u_2^2 e^{-t/\tau_2} + c_3 u_2^3 e^{-t/\tau_3} + q_2^{p_1} + q_2^{p_2} e^{-t/\tau_F}, \quad [110]$$

$$\tilde{l}_{a2}(t) = c_1 u_3^1 e^{-t/\tau_1} + c_2 u_3^2 e^{-t/\tau_2} + c_3 u_3^3 e^{-t/\tau_3} + q_3^{p_1} + q_3^{p_2} e^{-t/\tau_F}, \quad [111]$$

where $-1/\tau_1 = \zeta_1$, $-1/\tau_2 = \zeta_2$, and $-1/\tau_3 = \zeta_3$ with $\tau_1$, $\tau_2$, and $\tau_3$ denoting mechanical rise, fast adaptation, and slow adaptation time constants derived from the system. In matrix-vector format, this is given by

$$\boldsymbol{q}(t) = \mathbf{V} e^{\mathbf{J}t} \mathbf{c} + \boldsymbol{q}^{p_1} + \boldsymbol{q}^{p_2} e^{-t/\tau_F} \quad [112]$$

The remaining unknowns are the constants $c_1$, $c_2$, and $c_3$. We use the initial conditions $\tilde{\phi}(0) = \tilde{l}_{a1}(0) = \tilde{l}_{a2}(0) = 0$ to evaluate these unknowns to obtain their expressions as shown in equation 114.

$$\mathbf{0} = \mathbf{V} \begin{bmatrix} e^{-t/\tau_1} & 0 & 0 \\ 0 & e^{-t/\tau_2} & 0 \\ 0 & 0 & e^{-t/\tau_3} \end{bmatrix}_{t=0} \begin{pmatrix} c_1 \\ c_2 \\ c_3 \end{pmatrix} + \boldsymbol{q}^{p_1} + \boldsymbol{q}^{p_2} = \mathbf{V}\mathbf{c} + \boldsymbol{q}^{p_1} + \boldsymbol{q}^{p_2} \quad [113]$$

$$\mathbf{c} = -\mathbf{V}^{-1}(\boldsymbol{q}^{p_1} + \boldsymbol{q}^{p_2}) \quad [114]$$

Next, we use the linear solutions in equation 109-111 and equation 114 to obtain linearized bundle displacement as

$$X_{hb}^{lin} = r_{CA} \tilde{\phi}(t), \quad [115]$$

more elaborately given by

$$X_{hb}^{lin} = \sum_{i=1}^{3} \xi_i e^{-t/\tau_i} + \xi_F e^{-t/\tau_F} + \xi_F^{ss}, \quad [116]$$

where

$$\begin{aligned} \xi_1 &= r_{CA} c_1 u_1^1, \\ \xi_2 &= r_{CA} c_2 u_1^2, \\ \xi_3 &= r_{CA} c_3 u_1^3, \\ \xi_F &= r_{CA} q_1^{p_1}, \\ \xi_F^{ss} &= r_{CA} q_1^{p_2} \end{aligned} \quad [117]$$

and $\xi_1$, $\xi_2$, and $\xi_3$ are termed displacement response coefficients. Using first-order Taylor's expansion, we linearize the MET current shown in equation 43, obtaining the expression shown in equation 118.

$$I_{met}^{lin} = G_{max}(dV_{hb}^o - EP) \left[ \frac{\partial P_{o1}}{\partial \phi}\bigg|_e \tilde{\phi} + \frac{\partial P_{o1}}{\partial l_{a1}}\bigg|_e \tilde{l}_{a1} + \frac{\partial P_{o2}}{\partial \phi}\bigg|_e \tilde{\phi} + \frac{\partial P_{o2}}{\partial l_{a2}}\bigg|_e \tilde{l}_{a2} \right] \quad [118]$$

Substituting equations 109-111 in equation 118, we obtain the MET current as

$$I_{met}^{lin} = \sum_{i=1}^{3} \beta_i e^{-t/\tau_i} + \beta_F e^{-t/\tau_F} + \beta_F^{ss}, \quad [119]$$

where

$$\begin{aligned} \beta_1 &= G_{max}(dV_{hb}^o - EP) c_1 \left[ \frac{\partial P_{o1}}{\partial \phi}\bigg|_e u_1^1 + \frac{\partial P_{o2}}{\partial \phi}\bigg|_e u_1^1 + \frac{\partial P_{o1}}{\partial l_{a1}}\bigg|_e u_2^1 + \frac{\partial P_{o2}}{\partial l_{a2}}\bigg|_e u_3^1 \right], \\ \beta_2 &= G_{max}(dV_{hb}^o - EP) c_2 \left[ \frac{\partial P_{o1}}{\partial \phi}\bigg|_e u_1^2 + \frac{\partial P_{o2}}{\partial \phi}\bigg|_e u_1^2 + \frac{\partial P_{o1}}{\partial l_{a1}}\bigg|_e u_2^2 + \frac{\partial P_{o2}}{\partial l_{a2}}\bigg|_e u_3^2 \right], \\ \beta_3 &= G_{max}(dV_{hb}^o - EP) c_3 \left[ \frac{\partial P_{o1}}{\partial \phi}\bigg|_e u_1^3 + \frac{\partial P_{o2}}{\partial \phi}\bigg|_e u_1^3 + \frac{\partial P_{o1}}{\partial l_{a1}}\bigg|_e u_2^3 + \frac{\partial P_{o2}}{\partial l_{a2}}\bigg|_e u_3^3 \right], \\ \beta_F &= G_{max}(dV_{hb}^o - EP) \left[ \frac{\partial P_{o1}}{\partial \phi}\bigg|_e q_1^{p_2} + \frac{\partial P_{o2}}{\partial \phi}\bigg|_e q_1^{p_2} + \frac{\partial P_{o1}}{\partial l_{a1}}\bigg|_e q_2^{p_2} + \frac{\partial P_{o2}}{\partial l_{a2}}\bigg|_e q_3^{p_2} \right], \\ \beta_F^{ss} &= G_{max}(dV_{hb}^o - EP) \left[ \frac{\partial P_{o1}}{\partial \phi}\bigg|_e q_1^{p_1} + \frac{\partial P_{o2}}{\partial \phi}\bigg|_e q_1^{p_1} + \frac{\partial P_{o1}}{\partial l_{a1}}\bigg|_e q_2^{p_1} + \frac{\partial P_{o2}}{\partial l_{a2}}\bigg|_e q_3^{p_1} \right] \end{aligned} \quad [120]$$

and $\beta_1$, $\beta_2$, and $\beta_3$ are MET current response coefficients.

**F. Separating the fast and slow adaptation time constants.** The key to achieving both fast and slow adaptation dynamics in our three-row model lies in designing disparate system time constants that govern adaptation. To accomplish this, we applied a displacement clamp to the tallest stereocilia. In simpler terms, we used a displacement input model equivalent to an infinite probe stiffness model, which implies that $X_{hb}^{lin}(t) = x_{ext}^{base}(t)$ and that $\phi(t) = \sin^{-1}(X_{hb}(t)/r_{CA})$ is externally controlled. As a result, the linearized system is simplified to a two-degree-of-freedom system. In this context, the reduced-order Jacobian, $\mathcal{J}^{RO}$, is represented by the lower-right $2 \times 2$ block of $\mathcal{J}$ from equation 95. The reduced-order Jacobian $\mathcal{J}^{RO}$ is detailed in equation 121.

$$\mathcal{J}^{RO} = \begin{bmatrix} \frac{\partial}{\partial l_{a1}}(g_1 + g_2) & 0 \\ \frac{\partial m_3}{\partial l_{a1}} & \frac{\partial}{\partial l_{a2}}(m_1 + m_2) \end{bmatrix}_e \quad [121]$$

Evaluating the partial derivatives yields equation 122.

$$\mathcal{J}^{RO} = \begin{bmatrix} \mathcal{J}_{11}^{RO} & 0 \\ \mathcal{J}_{21}^{RO} & \mathcal{J}_{22}^{RO} \end{bmatrix}, \quad [122]$$

where

$$\mathcal{J}_{11}^{RO} = \frac{K_{GS_1}\epsilon_5^e}{\lambda_{a1}}\left[1 - \delta_1 d_1 P_{o1}^e(1 - P_{o1}^e)\right]\cos(\alpha_3^e) - \frac{K_{GS_1}\epsilon_6^e}{\lambda_{a1}}\left[\delta l_1^e - d_1 P_{o1}^e\right]\sin(\alpha_3^e) - \frac{K_{ES_1}}{\lambda_{a1}}, \quad [123]$$

$$\mathcal{J}_{21}^{RO} = \frac{\delta_1}{\lambda_{a1}} SF_{max_2}\epsilon_5 P_{o1}^e(1 - P_{o1}^e), \quad [124]$$

$$\mathcal{J}_{22}^{RO} = \frac{K_{GS_2}\epsilon_{11}^e}{\lambda_{a2}}\left[1 - \delta_2 d_2 P_{o2}^e(1 - P_{o2}^e)\right]\cos(\alpha_6^e) - \frac{K_{GS_2}\epsilon_{12}^e}{\lambda_{a2}}\left[\delta l_2^e - d_2 P_{o2}^e\right]\sin(\alpha_6^e) - \frac{K_{ES_2}}{\lambda_{a2}}, \quad [125]$$

where $\delta_1 = K_{GS_1} d_1/Nk_BT$, $\delta_2 = K_{GS_2} d_2/Nk_BT$, superscript "$e$" denotes the value at equilibrium achieved by applying the biasing displacement input, $P_{o1}^e = P_{o1}(\phi^e, l_{a1}^e)$, $P_{o2}^e = P_{o2}(\phi^e, l_{a2}^e)$, and $\phi^e$ is the row 1 rotation corresponding to the biasing displacement input. We can analytically compute the eigenvalues ($\zeta^{RO}$) for this reduced-order system and obtain approximate expressions for the time constants $\tau_2$ and $\tau_3$. The characteristic equation is given in equation 126.

$$\det(\mathcal{J}^{RO} - \zeta^{RO} I) = 0 \quad [126]$$

Equation 126 provides two eigenvalues given in equation 127 and 128.

$$\zeta_1^{RO} = \frac{K_{GS_1}\epsilon_5^e}{\lambda_{a1}}\left[1 - \delta_1 d_1 P_{o1}^e(1 - P_{o1}^e)\right]\cos(\alpha_3^e) - \frac{K_{GS_1}\epsilon_6^e}{\lambda_{a1}}\left[\delta l_1^e - d_1 P_{o1}^e\right]\sin(\alpha_3^e) - \frac{K_{ES_1}}{\lambda_{a1}} \quad [127]$$

$$\zeta_2^{RO} = \frac{K_{GS_2}\epsilon_{11}^e}{\lambda_{a2}}\left[1 - \delta_2 d_2 P_{o2}^e(1 - P_{o2}^e)\right]\cos(\alpha_6^e) - \frac{K_{GS_2}\epsilon_{12}^e}{\lambda_{a2}}\left[\delta l_2^e - d_2 P_{o2}^e\right]\sin(\alpha_6^e) - \frac{K_{ES_2}}{\lambda_{a2}} \quad [128]$$

Now, $\tau_2$ can be related to $-1/\zeta_1^{RO}$ and $\tau_3$ to $-1/\zeta_2^{RO}$. Note that in the $3 \times 3$ system, these time constants would be slightly different, but they would still depend on the parameters in equations 127 and 128. Therefore, at equilibrium, we obtain the approximate explicit expressions for $\tau_2$ and $\tau_3$ as shown in equations 129 and 130.

$$\tau_2 \approx \frac{\lambda_{a1}}{K_{ES_1} - K_{GS_1}\left\{\epsilon_5^e\left[1 - \delta_1 d_1 P_{o1}^e(1 - P_{o1}^e)\right]\cos(\alpha_3^e) - \epsilon_6^e\left[\delta l_1^e - d_1 P_{o1}^e\right]\sin(\alpha_3^e)\right\}} \quad [129]$$

$$\tau_3 \approx \frac{\lambda_{a2}}{K_{ES_2} - K_{GS_2}\left\{\epsilon_{11}^e\left[1 - \delta_2 d_2 P_{o2}^e(1 - P_{o2}^e)\right]\cos(\alpha_6^e) - \epsilon_{12}^e\left[\delta l_2^e - d_2 P_{o2}^e\right]\sin(\alpha_6^e)\right\}} \quad [130]$$

It is important to note that, up to this point, there is no distinction that identifies $\tau_2$ as the fast adaptation time constant and $\tau_3$ as the slow adaptation time constant. However, equations 129 and 130 indicate which parameters can be adjusted to distinguish between the two time constants, with one being on the order of 0.1 ms and the other on the order of tens of milliseconds.

Studies have shown that fast adaptation may not depend on calcium (8, 9), while ample evidence indicates that slow adaptation does rely on calcium (1, 2, 10). In our model, the adaptation complex in the tallest row is calcium-independent (see equation 27) due to the slow ion diffusion to row 1 UTLD, whereas the middle-row adaptation complex is calcium-dependent (see equation 28) due to its proximity to the row 2 LTLD. Therefore, the following design is based on experimental observations: the tallest row adapts to account for fast adaptation dynamics, and the middle row adapts to contribute to the slow adaptation of the MET current. Consequently, $\tau_2$ represents the fast adaptation time constant, and $\tau_3$ represents the slow adaptation time constant in our model.

To determine $\tau_2$ and $\tau_3$ in equations 129 and 130, respectively, we differentiate the parameters $K_{ES_1}$ and $K_{ES_2}$, as well as $\lambda_{a1}$ and $\lambda_{a2}$. To simplify the model and reduce the number of parameters, we assume equality among similar elements: $K_{GS} = K_{GS_1} = K_{GS_2}$, $K_{SP} = K_{SP_1} = K_{SP_2} = K_{SP_3}$, $d = d_1 = d_2$, $\lambda = \lambda_1 = \lambda_2 = \lambda_3$, and $l^{act} = l_1^{act} = l_2^{act}$. By varying the extent spring stiffnesses and the intracellular damping coefficients, we distinguish between these time constants. To achieve a smaller value for $\tau_2$, we impose the constraints $\lambda_{a1} < \lambda_{a2}$ and $K_{ES_1} > K_{ES_2}$ (refer to Table S1). This approach helps clearly separate the two time constants.

### 3. The Three-Row Model with Linearized Geometry

In this section, we derive a simpler version of our three-row model described in Section 1 with linearized geometry and retained constitutive nonlinearity to gauge the significance of geometric nonlinearity in bundle dynamics. We commence by linearizing the geometry and subsequently re-deriving the EOM. A version of this model with only two rows is elaborated upon in our previous work (11). The geometry is linearized about the vertical bundle position ($\phi = l_{a1} = l_{a2} = 0$). The linearized geometric expressions are shown in equations 131-138.

$$l_1 \approx l_1^o + \epsilon_4^o\tilde{\phi} + \epsilon_5^o\tilde{l}_{a1}, \quad [131]$$

$$l_2 \approx l_2^o + \epsilon_{10}^o\tilde{\phi} + \epsilon_{11}^o\tilde{l}_{a2}, \quad [132]$$

$$\alpha_3 \approx \alpha_3^o + \epsilon_3^o\tilde{\phi} + \epsilon_6^o\tilde{l}_{a1}, \quad [133]$$

$$\alpha_6 \approx \alpha_6^o + \epsilon_9^o\tilde{\phi} + \epsilon_{12}^o\tilde{l}_{a2}, \quad [134]$$

$$\begin{aligned} l_1^m \approx l_1^{m,lin} &= l_1^{m,o} + [l_1^{gs}\cos(\alpha_3^o) - r_1\sin(\alpha_3^o)]\,\epsilon_3^o\tilde{\phi} + [(l_1^{gs}\cos(\alpha_3^o) - r_1\sin(\alpha_3^o))\epsilon_6^o - \sin(\alpha_3^o)]\,\tilde{l}_{a1} \\ &= l_1^{m,o} + \epsilon_{13}^o\tilde{\phi} + \epsilon_{14}^o\tilde{l}_{a1}, \end{aligned} \quad [135]$$

$$\begin{aligned} l_2^m \approx l_2^{m,lin} &= l_2^{m,o} + [(l_2^{gs}\cos(\alpha_6^o) - r_2\sin(\alpha_6^o))\epsilon_9^o - b_{12}\sin(\alpha_2^o - \alpha_6^o)(\epsilon_1^o - \epsilon_9^o)]\,\tilde{\phi} \\ &+ [(l_2^{gs}\cos(\alpha_6^o) - r_2\sin(\alpha_6^o))\epsilon_{12}^o + b_{12}\sin(\alpha_2^o - \alpha_6^o)\epsilon_{12}^o - \sin(\alpha_6^o)]\,\tilde{l}_{a2} \\ &= l_1^{m,o} + \epsilon_{15}^o\tilde{\phi} + \epsilon_{16}^o\tilde{l}_{a2}, \end{aligned} \quad [136]$$

$$\cos(\alpha_3) \approx c_1^{lin} = \cos(\alpha_3^o) - \sin(\alpha_3^o)\epsilon_3^o\tilde{\phi} - \sin(\alpha_3^o)\epsilon_6^o\tilde{l}_{a1} = \cos(\alpha_3^o) + \epsilon_{17}^o\tilde{\phi} + \epsilon_{18}^o\tilde{l}_{a1}, \quad [137]$$

$$\cos(\alpha_6) \approx c_2^{lin} = \cos(\alpha_6^o) - \sin(\alpha_6^o)\epsilon_9^o\tilde{\phi} - \sin(\alpha_6^o)\epsilon_{12}^o\tilde{l}_{a2} = \cos(\alpha_6^o) + \epsilon_{19}^o\tilde{\phi} + \epsilon_{20}^o\tilde{l}_{a2}, \quad [138]$$

where the superscript "$o$" denotes the value when $\phi = l_{a1} = l_{a2} = 0$ and overhead ($\sim$) denotes linearized coordinates (for instance, $\tilde{\phi} = \phi - \phi^o$).

**A. Force input.** After linearizing only the geometric relations, we use Lagrange's energy method (5) to re-derive the EOM (as was described in Section 2G) with linearized geometry yielding (neglecting higher-order terms $\geq 2$),

$$\begin{aligned} \Lambda^o\frac{d\tilde{\phi}}{dt} &= -K_{GS_1}(\epsilon_4^o l_1^{m,o}\tilde{\phi} + \epsilon_5^o l_1^{m,o}\tilde{l}_{a1} - d_1 l_1^{m,lin}P_{o1}) - K_{GS_2}(\epsilon_{10}^o l_2^{m,o}\tilde{\phi} + \epsilon_{11}^o l_2^{m,o}\tilde{l}_{a2} - d_2 l_2^{m,lin}P_{o2}) \\ &- \left[K_{SP_1} + (\epsilon_1^o)^2K_{SP_2} + (\epsilon_7^o)^2K_{SP_3}\right]\tilde{\phi} + F_{ext}(t)r_{CA}\cos(\tilde{\phi}), \end{aligned} \quad [139]$$

$$\lambda_{a1}\frac{d\tilde{l}_{a1}}{dt} = K_{GS_1}(\epsilon_4^o\cos(\alpha_3^o)\tilde{\phi} + \epsilon_5^o\cos(\alpha_3^o)\tilde{l}_{a1} - d_1c_1^{lin}P_{o1}) - K_{ES_1}\tilde{l}_{a1} - F_{max_1}, \quad [140]$$

$$\lambda_{a2}\frac{d\tilde{l}_{a2}}{dt} = K_{GS_2}(\epsilon_{10}^o\cos(\alpha_6^o)\tilde{\phi} + \epsilon_{11}^o\cos(\alpha_6^o)\tilde{l}_{a2} - d_2c_2^{lin}P_{o2}) - K_{ES_2}\tilde{l}_{a2} - F_{max_2}(1 - SP_{o1}), \quad [141]$$

where $\Lambda^o = \lambda_1 + (\epsilon_1^o)^2\lambda_2 + (\epsilon_7^o)^2\lambda_3$, channel open probabilities

$$P_{o1} = \frac{1}{1 + \exp\left[-\frac{K_{GS_1}d_1}{Nk_BT}(\epsilon_4^o\tilde{\phi} + \epsilon_5^o\tilde{l}_{a1} - l_1^{act})\right]}, \quad [142]$$

$$P_{o2} = \frac{1}{1 + \exp\left[-\frac{K_{GS_2}d_2}{Nk_BT}(\epsilon_{10}^o\tilde{\phi} + \epsilon_{11}^o\tilde{l}_{a2} - l_2^{act})\right]}, \quad [143]$$

and $\epsilon_k$ $\forall k \in [1, 12]$ and $k \in \mathcal{Z}$ are defined in Table S2.

**B. Stiff probe displacement input.** In this case, we follow the same procedure as in Section 3A to obtain partially linearized model with a displacement input shown in equation 144-146.

$$\Lambda^o \frac{d\tilde{\phi}}{dt} = -K_{GS_1}(\epsilon_4^o l_1^{m,o}\tilde{\phi} + \epsilon_5^o l_1^{m,o}\tilde{l}_{a1} - d_1 l_1^{m,lin}P_{o1}) - K_{GS_2}(\epsilon_{10}^o l_2^{m,o}\tilde{\phi} + \epsilon_{11}^o l_2^{m,o}\tilde{l}_{a2} - d_2 l_2^{m,lin}P_{o2})$$
$$- \left[K_{SP_1} + (\epsilon_1^o)^2 K_{SP_2} + (\epsilon_7^o)^2 K_{SP_3}\right]\tilde{\phi} - K_P r_{CA}^2 \cos(\tilde{\phi})\sin(\tilde{\phi}) + K_P X_{ext}^{base}(t) r_{CA}\cos(\tilde{\phi}), \quad [144]$$

$$\lambda_{a1}\frac{d\tilde{l}_{a1}}{dt} = K_{GS_1}(\epsilon_4^o \cos(\alpha_3^o)\tilde{\phi} + \epsilon_5^o \cos(\alpha_3^o)\tilde{l}_{a1} - d_1 c_1^{lin}P_{o1}) - K_{ES_1}\tilde{l}_{a1} - F_{max_1}, \quad [145]$$

$$\lambda_{a2}\frac{d\tilde{l}_{a2}}{dt} = K_{GS_2}(\epsilon_{10}^o \cos(\alpha_6^o)\tilde{\phi} + \epsilon_{11}^o \cos(\alpha_6^o)\tilde{l}_{a2} - d_2 c_2^{lin}P_{o2}) - K_{ES_2}\tilde{l}_{a2} - F_{max_2}(1 - SP_{o1}), \quad [146]$$

**C. Model outputs.** Finally, the motion of the bundle along the $x$-axis is given by

$$X_{hb}^{glin} = r_{CA}\sin(\tilde{\phi}), \quad [147]$$

and the MET current from this model is given by

$$I_{met}^{glin} = \frac{G_{max}(dV_{hb}^o - EP)}{1 + \exp\left[-\frac{K_{GS_1}d_1}{Nk_BT}(\epsilon_4^o\tilde{\phi} + \epsilon_5^o\tilde{l}_{a1} - l_1^{act})\right]} + \frac{G_{max}(dV_{hb}^o - EP)}{1 + \exp\left[-\frac{K_{GS_2}d_2}{Nk_BT}(\epsilon_{10}^o\tilde{\phi} + \epsilon_{11}^o\tilde{l}_{a2} - l_2^{act})\right]}, \quad [148]$$

where the mechanical and electrical parameter definitions and values are tabulated in Table S1.

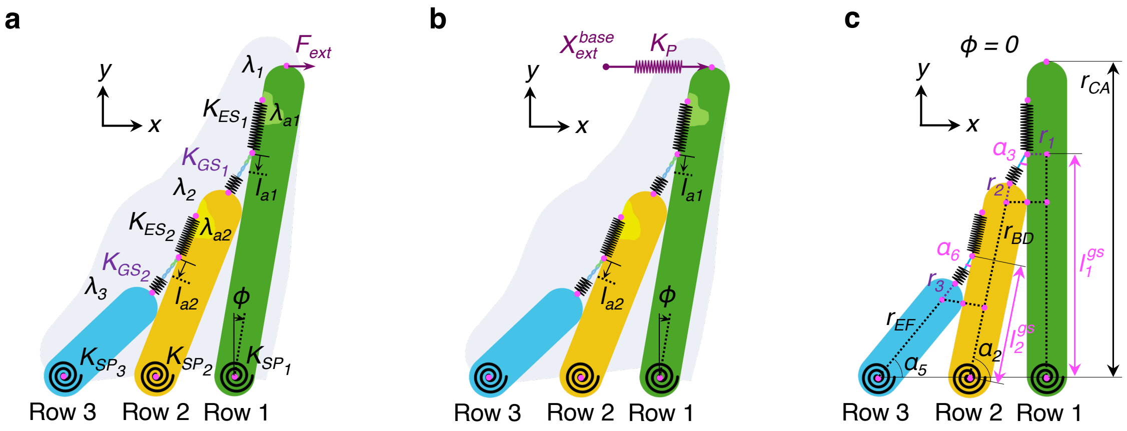


**Fig. S1.** Description of tip link densities and schematic of the three-row model. **a** Illustrates the dynamic three-row model, highlighting the various viscoelastic components: $K_{GS_j}$, $K_{SP_i}$, $K_{ES_j}$, $\lambda_i$, and $\lambda_{aj}$ for $i \in [1, 3]$ and $j \in [1, 2]$, as defined in Table S1. The model considers three degrees of freedom: bundle rotation ($\phi$) and the displacements of the two adaptation complexes ($l_{a1}$ and $l_{a2}$). **b** Demonstrates the same three-row model under the influence of a stiff probe. The probe, depicted as a spring with stiffness $K_P$, is driven by a displacement input at its free end. This setup illustrates how external mechanical stimulation is applied to the system. **c** Defines the important geometric lengths and angles relevant to the model when bundle rotation ($\phi$) is zero. Length estimates, such as the total length of row 2 ($r_{BD} + r_2$) and row 3 ($r_{EF} + r_3$), are derived from (6). The geometric parameter values are listed in Table S1. Note that the schematic is not drawn to scale concerning these values.

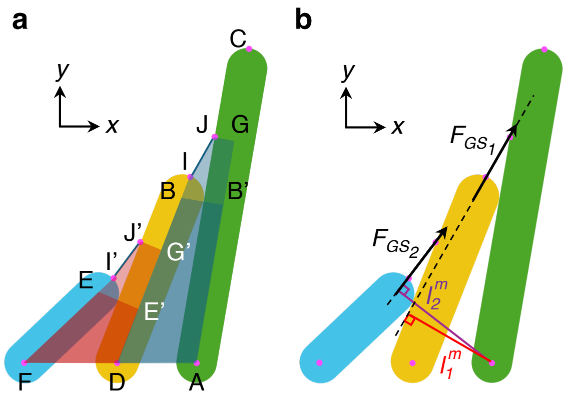


**Fig. S2.** Polygons used for deriving geometric relations and a description of moment arms. **a** The polygons used to evaluate the relation between time-varying unknowns in terms of known geometric parameters, and the three generalized coordinates are $AB'GJIBD$, $AB'BD$, $DE'G'J'I'EF$, and $DE'EF$. **b** Describes the moment-arm lengths $l_1^m$ and $l_2^m$ for the two gating spring tensions $F_{GS_1}$ and $F_{GS_2}$, respectively, from the tallest stereocilium's pivot point $A$. The lever arms depend on $\phi$, $l_{a1}$, and $l_{a2}$.

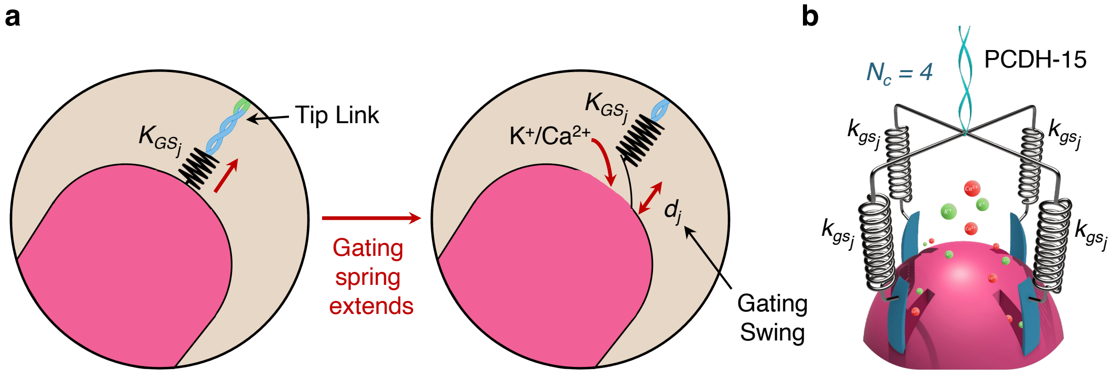


**Fig. S3.** Gating mechanism at the LTLD. **a** Illustration of the state change from a closed gate (left) to an open gate (right) as gating spring tension increases, allowing cation influx into the stereocilia. Each gate has an associated gating swing ($d_1$ or $d_2$) that limits its operational extent. **b** Depiction of multiple channels. Each tip link (specifically protocadherin-15 (PCDH-15) at the LTLD (12)) pulls a nexus of $N_c$ gates through $N_c$ gating springs operating in parallel ($k_{gs_j} = \kappa_{GS_j,single}/N_c$). The illustration shows $N_c = 4$ with calcium ions (red balls) and potassium ions (green balls) diffusing into the channels. In our model, $\kappa_{GS_j,single}$ denotes the total gating spring stiffness of a single stereocilium between rows 1 and 2 ($j = 1$) or rows 2 and 3 ($j = 2$). However, we homogenize the multi-column HB into a single column. Therefore, in our model with 30 columns, $K_{GS_j} = 30\kappa_{GS_j,single}$. We do not explicitly define $\kappa_{GS_j,single}$ in Table S1 as it is recognized as an intermediate parameter.

**Table S1. Model parameters, their description, and values.**

| Parameter | Description | Value | Units |
|---|---|---|---|
| $K_{SP_i}$* | Combined torsional pivot stiffness of the $i^{th}$ stereocilium across all columns | 15 | fN.m/rad |
| $K_{GS_j}$* | Stiffness of the gating spring between $j^{th}$ and $(j+1)^{th}$ stereocilia summed across all columns in the HB | 145 | mN/m |
| $K_{ES_1}$ | Stiffness of the extent spring associated with the adaptation complex in the tallest stereocilia UTLD across all HB columns | 50 | mN/m |
| $K_{ES_2}$ | Stiffness of the extent spring associated with the adaptation complex in the middle stereocilia UTLD across all HB columns | 25 | mN/m |
| $K_P$ | Probe stiffness in the displacement-stimulated model | 200 | mN/m |
| $\lambda_i$* | Sum of the damping coefficients (representing viscous and shear effects) in the extracellular vicinity of the $i^{th}$ stereocilium across all columns | 16.13 | aN.m.s/rad |
| $\lambda_{a1}$ | Intracellular damping coefficient (viscous and shear effects) in the vicinity of the tallest stereocilia adaptation complex combined for all columns | 18 | $\mu$N.s/m |
| $\lambda_{a2}$ | Intracellular damping coefficient (viscous and shear effects) in the vicinity of the middle stereocilia adaptation complex combined for all columns | 960 | $\mu$N.s/m |
| $d_j$* | Gating swing for the gate between $j^{th}$ and $(j+1)^{th}$ stereocilia | 2.77 | nm |
| $N_s$ | Total number of stereocilia in the HB | 90 (3, 7) | |
| $N_{TL}$ | Number of tip links in the HB | 60 (3, 7) | |
| $N_c$ | Number of transduction channels per tip link | 4 (3) | |
| $N$ | Combined transduction channels for each row pair in the HB ($= N_c N_{TL}/2$) | 120 | |
| $k_B$ | Boltzmann constant | $1.38 \times 10^{-23}$ | $m^2.kg.s^{-2}.K^{-1}$ |
| $T$ | Cell temperature | 300 | K |
| $F_{max_1}$ | Maximum stall force of tallest stereocilia UTLD summed across all columns | 10 | pN |
| $F_{max_2}$ | Maximum stall force of middle stereocilia UTLD summed across all columns | 40 | pN |
| $l_j^{act*}$ | Resting extension of $j^{th}$ gating spring | 26.45 | pm |
| $S$ | Instantaneous feedback strength of cations on the adaptation complex | 1 | |
| $G_{max}$ | Maximum conductance of the transduction channels at each LTLD combined for all columns | 5.04 | nS |
| $\Delta V_{hb}^o$ | Steady-state holding potential across the apical cell membrane of the OHC | -80 (1) | mV |
| $EP$ | Endo-cochlear potential at rest | 0 | mV |
| $r_1$ | Radius of the tallest stereocilia (row 1) | 113 (6) | nm |
| $r_2$ | Radius of the middle stereocilia (row 2) | 118 (6) | nm |
| $r_3$ | Radius of the shortest stereocilia (row 3) | 82.45 (6) | nm |
| $r_{CA}$ | Height of the tallest stereocilia (row 1) | 4.2 (6) | $\mu$m |
| $r_{BD} + r_2$ | Height of the middle stereocilia (row 2) | 2.2 (6) | $\mu$m |
| $r_{EF} + r_3$ | Height of the shortest stereocilia (row 3) | 1.1 (6) | $\mu$m |
| $b_{12}$ | Horizontal separation between rows 1 and 2 | 0.61 (6) | $\mu$m |
| $b_{23}$ | Horizontal separation between rows 2 and 3 | 0.61 (6) | $\mu$m |
| $l_1^{gs}$ | Height of the tip link insertion in row 1 from its pivot point when $\phi = 0$ | 2.3 (6) | $\mu$m |
| $l_2^{gs}$ | Height of the tip link insertion in row 2 from its pivot point when $\phi = 0$ | 1.0 (6) | $\mu$m |

* $i \in [1,3]$ and $j \in [1,2]$. Both $i$ and $j$ are integers.

**Table S2. Definitions of partial derivatives for the three-row model.**

| Rows 1 and 2 | | Rows 2 and 3 | |
|---|---|---|---|
| Partial Derivative | Variable | Partial Derivative | Variable |
| $\partial\alpha_2/\partial\phi$ | $\epsilon_1$ | $\partial\alpha_5/\partial\phi$ | $\epsilon_7$ |
| $\partial r_{B'A}/\partial\phi$ | $\epsilon_2$ | $\partial r_{E'D}/\partial\phi$ | $\epsilon_8$ |
| $\partial\alpha_3/\partial\phi$ | $\epsilon_3$ | $\partial\alpha_6/\partial\phi$ | $\epsilon_9$ |
| $\partial l_1/\partial\phi$ | $\epsilon_4$ | $\partial l_2/\partial\phi$ | $\epsilon_{10}$ |
| $\partial l_1/\partial l_{a1}$ | $\epsilon_5$ | $\partial l_2/\partial l_{a2}$ | $\epsilon_{11}$ |
| $\partial\alpha_3/\partial l_{a1}$ | $\epsilon_6$ | $\partial\alpha_6/\partial l_{a2}$ | $\epsilon_{12}$ |

**Table S3. Tuned parameters in the two-row model to match the fluid-jet experimental data.**

| Parameter | Value | Units |
|---|---|---|
| $K_{SP}$ | 1.1 | mN/m |
| $K_{GS}$ | 2 | mN/m |
| $K_{SP}$ | 2 | mN/m |
| $K_P$ | 200 | mN/m |
| $\lambda$ | 1.15 | $\mu$N.s/m |
| $\lambda_a$ | 240 | $\mu$N.s/m |
| $D$ | 19.8 | nm |
| $N$ | 120 | |
| $k_B$ | $1.38 \times 10^{-23}$ | $m^2.kg.s^{-2}.K^{-1}$ |
| $T$ | 300 | K |
| $F_{max}$ | 160 | pN |
| $S$ | 6 | |
| $G_{max}$ | 10.038 | nS |
| $\Delta V_{hb}^o$ | -80 (1) | mV |
| $EP$ | 0 | mV |

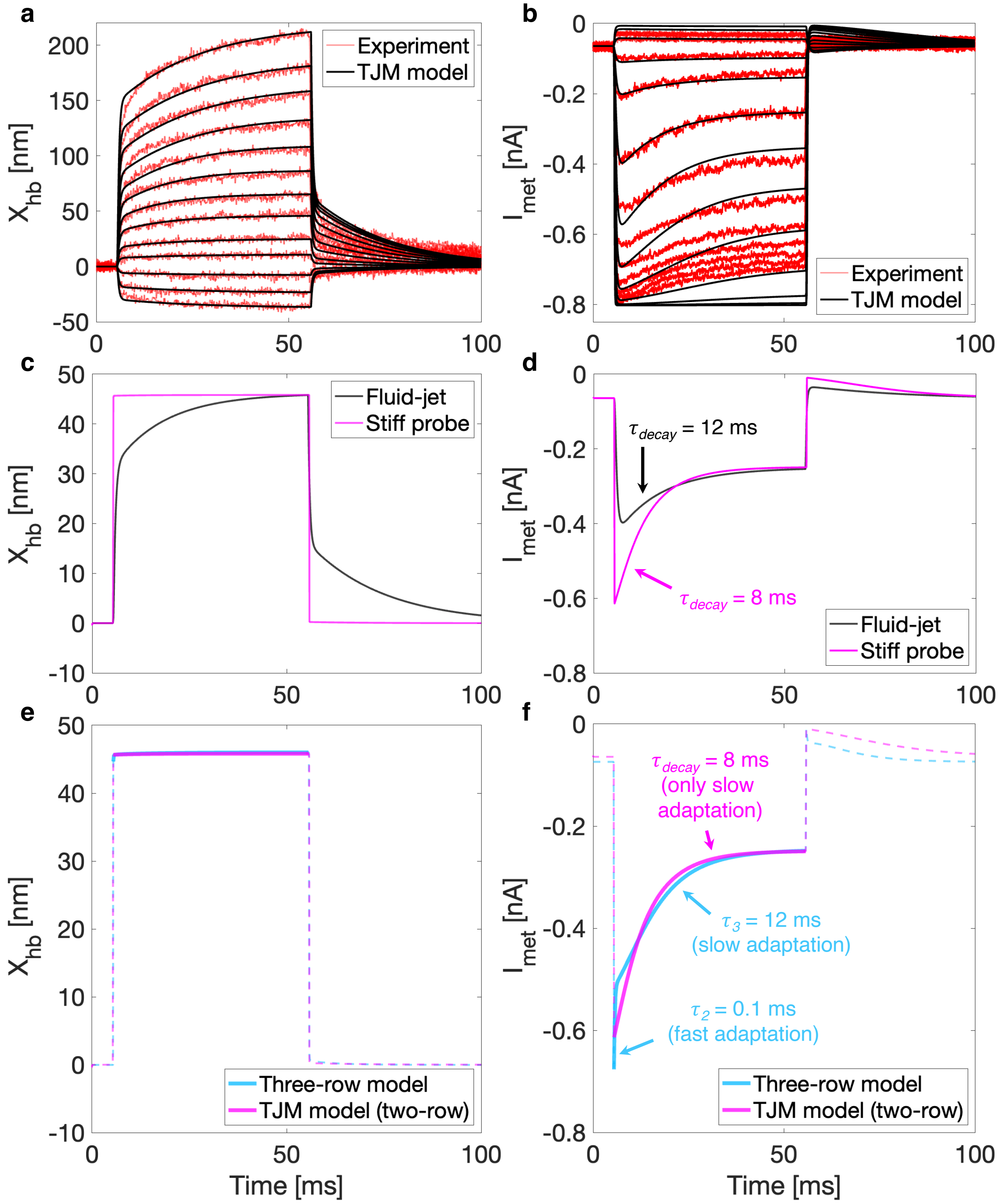


**Fig. S4.** Response predictions from a two-row model under fluid-jet and stiff probe stimulations and their comparison with experiments and our three-row model. **a-d** Results from the two-row model (4). **a** Experimental HB displacements (1) (red) compared to the two-row model (developed by Tinevez *et al.* (4), denoted the TJM model) predictions. The TJM model, having only two time constants, uses one to capture the mechanical rise and the other to govern creep. The model parameters are listed in Table S3. **b** Comparison between experimental (red) and TJM model (black) MET currents. Here, one time constant drives the mechanical rise, while the second drives slow adaptation. The force amplitudes used in the fluid-jet model are -90, -55, -17, 22, 48, 83, 113, 145, 180, 225, 285, 345, and 430 pN. **c** Comparison of TJM model HB displacements under fluid-jet (black) and stiff probe (magenta) stimulations. The stiff probe model does not display a creep. **d** Corresponding MET currents for fluid-jet (black) and stiff probe (magenta) stimulations in the TJM model show only slow decay due to the model's limitation of predicting only two time constants. The slow adaptation time constant was 12 ms for the fluid-jet model (decay fit: $0.14 - 0.14\exp(-0.0825t)$, $R^2 = 1$) and 8 ms for the stiff probe model (decay fit: $0.35 - 0.36\exp(-0.1239t)$, $R^2 = 0.9998$), which still falls under the slow regime. The fluid-jet force amplitude in the simulation was 83 pN, and the probe displacement amplitude was 46.23 nm. **e-f** Results from the TJM model for a 46.23 nm probe displacement are compared to results from our three-row model actuated by a 47 nm probe displacement. **e** Comparison of HB displacements between stiff probe TJM model simulations (magenta) and three-row model predictions (blue), showing displacement similarities between the two models. **f** MET currents for similar HB displacements from two-row (magenta) and three-row (blue) models. The three-row model, incorporating three time constants, captures both fast and slow adaptation, evidenced by a double exponential fit to the blue trace: $0.36 - 0.08\exp(-10.46t) - 0.28\exp(-0.0846t)$, $R^2 = 0.9977$.

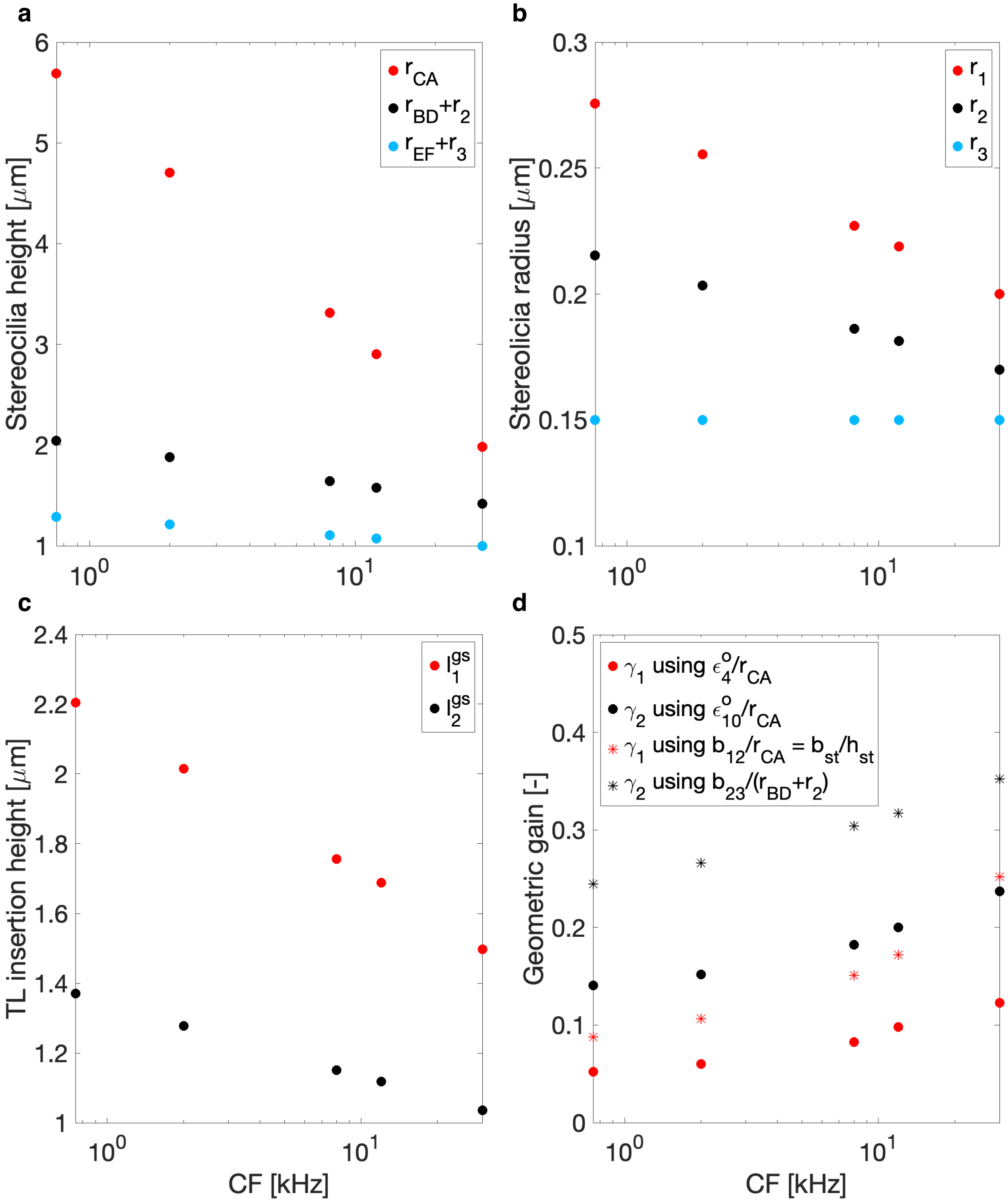


**Fig. S5.** Tonotopic variation of morphology and geometric gain from a mouse model. Tonotopic variation of **a** the heights of the three rows of stereocilia of an OHC HB, **b** the radii of the three rows of stereocilia of an OHC HB, **c** the assumed tip link (TL) insertion points in the model, and **d** the two geometric gains ($\gamma_1$ and $\gamma_2$) obtained from each adjacent row-pair using the traditional definition ($b_{st}/h_{st}$, shown by asterisks) and the exact expression ($\epsilon_4^o/r_{CA}$), obtained when the bundle is upright ($\phi = 0$, shown by solid circles). The data is shown for five characteristic frequency (CF) locations: 30 kHz, 12 kHz, 8 kHz, 2 kHz, and 750 Hz. The dimensions of the stereocilia in A and B are from the first row of OHCs (radially closest to the IHCs), as described in Yarin *et al.* (13) for adult mice. In our model, the heights of row 2 and row 3 are defined as $r_{BD} + r_2$ and $r_{EF} + r_3$, respectively, as detailed in Table S1. In **c**, we assume tip link (TL) insertion heights that ensure tip link lengths lie within the physiological range, consistent with previous studies (14–17). **d** shows that the conventional geometric gain expression ($b_{st}/h_{st}$) consistently overestimates the value compared to the exact gain expression ($\epsilon^o/r_{CA}$). Note that $b_{st} = 0.5$ $\mu$m (18) was assumed to be constant across all frequencies due to the unavailability of experimental tonotopic variation of the stereociliary separation. This estimate, however, lies within the experimental range found at specific cochlear locations from rat OHC (7), hamster IHC (19), and mice IHC (20).

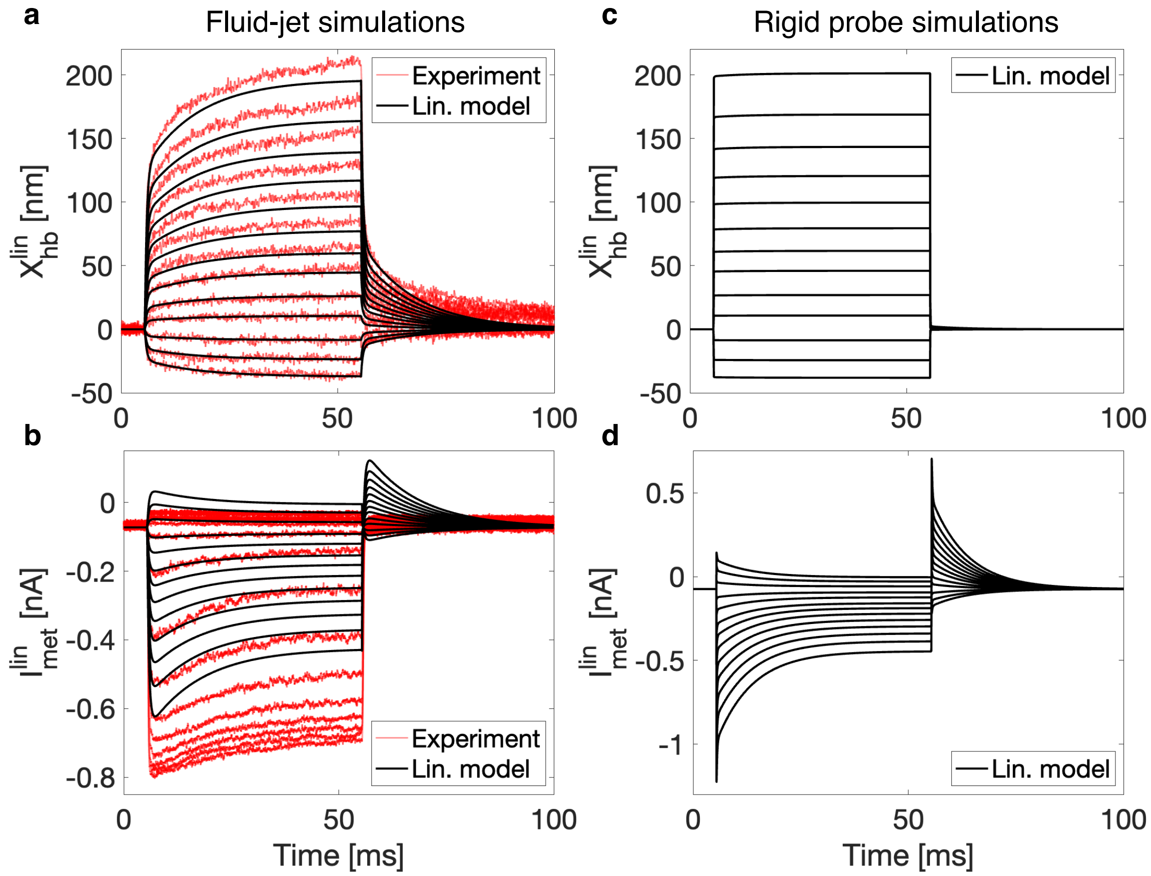


**Fig. S6.** Predictions of HB displacement and MET current responses using linearized three-row fluid-jet and stiff probe models. Panels A and B show response time courses for thirteen distinct stimulation amplitudes simulated with the fully linearized fluid-jet model. **a** HB displacement responses, showing close alignment with experimental data up to approximately 50 nm, where deviations begin, occurring earlier than predicted by the linearized geometry model (Section 3, Fig. S7). **b** MET current responses demonstrate quantitative agreement with experiments but show an overestimation of inhibitory forces and an underestimation of excitatory forces. The applied force amplitudes were: -180, -115, -41, 51, 127, 217, 291, 375, 470, 569, 677, 797, and 950 pN, consistent with the nonlinear model. Panels **c** and **d** show the HB displacement and MET current responses from the stiff probe model. **c** Displacements exhibit a rapid mechanical rise due to the added probe stiffness. **d** MET currents show biphasic adaptation but lack an upper bound of 0, as in the nonlinear model, due to the omission of higher-order terms in the linearization. Base displacements for these responses were: -39, -25, -9, 11, 27, 47, 63, 81, 102, 123, 147, 173, and 206 nm. In both models, MET currents do not saturate, and the magnitude of adaptation increases with higher force amplitudes.

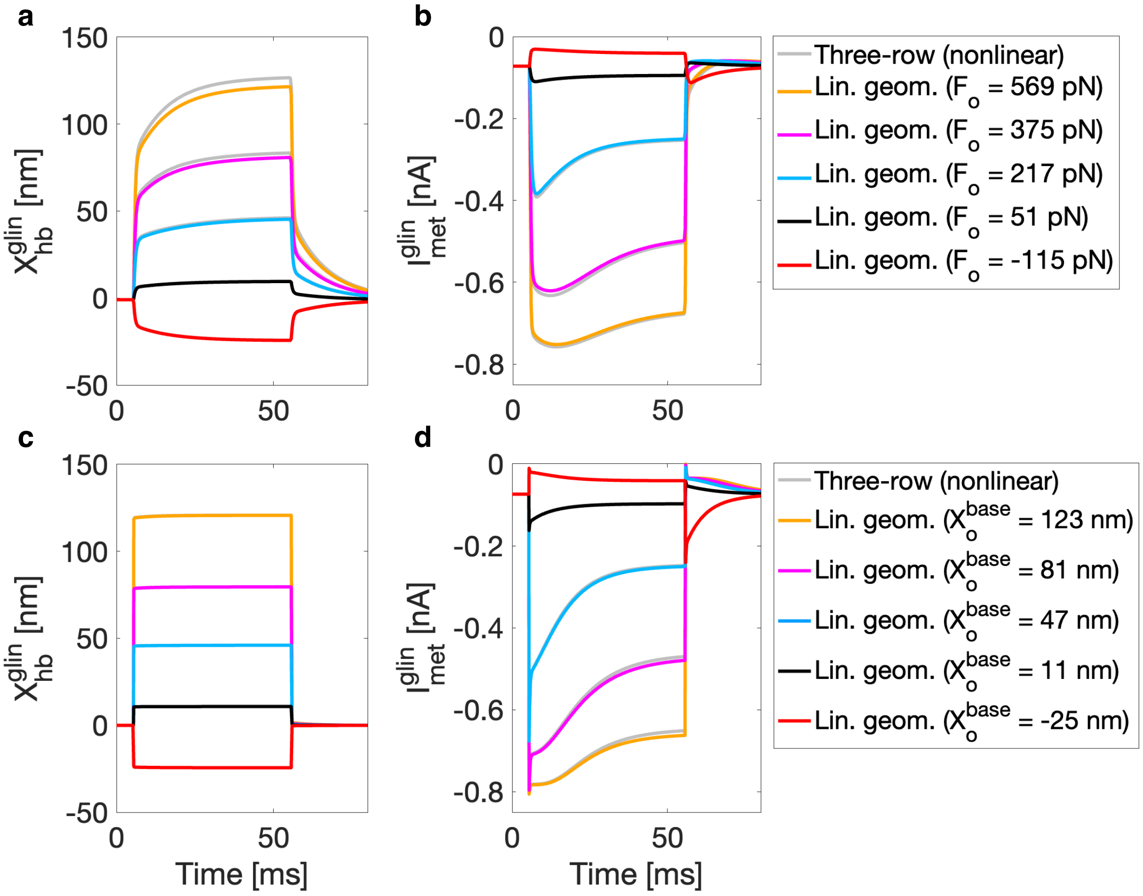


**Fig. S7.** Comparison of HB displacement and MET current responses between the three-row model with linearized geometry and the fully nonlinear three-row model under fluid-jet and stiff probe stimulations. **a** HB displacements from the fluid-jet model with linearized geometry for five force amplitudes (-115 pN (red), 51 pN (black), 217 pN (blue), 375 pN (magenta), and 569 pN (green)), compared with displacements from the fully nonlinear model. Linearization begins to underestimate displacements beyond 75 nm. **b** Corresponding MET currents from the fluid-jet model show slight underestimation at larger bundle displacements. **c** HB displacements from the stiff probe models are nearly identical due to the probe's large stiffness (see Table S1). Probe displacements are shown for -25 nm (red), 11 nm (black), 47 nm (blue), 81 nm (magenta), and 123 nm (green). **d** MET currents in the stiff probe model, similar to the fluid-jet model, start to diverge beyond ∼75 nm, though underestimation due to geometric linearization is minor compared to the fully nonlinear model.
